# Identification and characterization of bacterial repeat-in-toxin adhesins using long-read genome analysis

**DOI:** 10.1101/2025.09.30.679566

**Authors:** Thomas Hansen, Laurie A. Graham, Blake P. Soares, Daniel Lee, Justin R. Gagnon, Trina Dykstra-MacPherson, Shuaiqi Guo, Peter L. Davies

## Abstract

Gram-negative bacteria attach to host surfaces using ligand-binding domains (LBDs) at the distal tips of fibrillar RTX adhesins. Blocking the initial binding interaction(s) can potentially prevent colonization and subsequent biofilm formation and infection. To this end, adhesins must be identified, and it is also essential to determine the dominant adhesin type for those species that have more than one. RTX adhesins are frequently the largest proteins within each species (ranging from 1500 to 15,000 aa) and are often misannotated as incomplete/pseudogene products because their highly repetitive nature confounds genome assemblies from short-read technologies. Our bioinformatic process collates predicted proteins from long-read assemblies, which are then clustered based on the similarity of their C-terminal regions where the LBDs are typically located. RTX adhesins are identified by their length and domain structure and are modelled using AlphaFold3. An exhaustive search of multiple strains from seven species revealed a total of 35 different RTX adhesins that map to 16 different loci, with differing arrangements of LBDs that include putative carbohydrate-binding modules and von Willebrand Factor A-like domains. Notably, similar adhesins are sometimes found in multiple species, either by descent or through DNA uptake, and three species have an RTX adhesin of uncertain function because it lacks an obvious LBD.

**Importance:** Many bacteria initiate infection by reaching out with large, complex proteins called adhesins to attach themselves to a host cell. The DNA sequences of adhesins are difficult to read, due to their length and repetitiveness. This study leverages recent technological advances like “long-read sequencing” and structural modelling to identify and characterize the adhesins in seven species of harmful bacteria: *Acinetobacter baumannii*, *Aeromonas hydrophila*, *Aeromonas salmonicida*, *Bordetella parapertussis*, *Legionella pneumophila*, *Vibrio parahaemolyticus*, and *Vibrio vulnificus*. We identified thirty-five unique versions of adhesins and demonstrated their mix-and-match architecture. This research provides a foundation for strategies to block bacteria from binding surfaces, offering a vital alternative treatment as antibiotic resistance continues to rise.

## Introduction

Fibrillar adhesins (FAs) are long, single-chain bacterial surface proteins that fold as a string of domains ^1–3^. Growing evidence suggests that FAs make initial contact with various surfaces through ligand-binding domains (LBDs) found at their distal end ^4–8^. These initial contacts include abiotic surfaces ^9–11^, as well as the surfaces of host cells ^8, 12^. Once colonization is underway, FAs also contribute to the formation of biofilms, which exacerbate infections and shield the bacteria from both host defenses and antibiotics ^11, 13–21^.

Repeat-in-toxin (RTX) adhesins are a subset of FAs that are secreted through the type I secretion system (T1SS) ^22–24^, and a generic schematic of their domain architecture is shown in Fig. 1A. They have a T1SS signal at the C terminus preceded by a characteristic Ca^2+^-binding nonapeptide repeat sequence (from which the RTX name is derived ^24^) that folds into a β-roll/solenoid ^25^. It has been suggested that the Ca^2+^-dependent folding of this domain and those that emerge subsequently prevent reversal of secretion and may help extrude the adhesin into the Ca^2+^-rich milieu outside of the bacterium ^24, 26^. Only the very N-terminal retention module (RM) folds in the absence of Ca^2+^, which enables it to serve as a periplasmic plug to terminate secretion and anchor the adhesin to the cell surface ^27, 28^.

**Figure 1.**
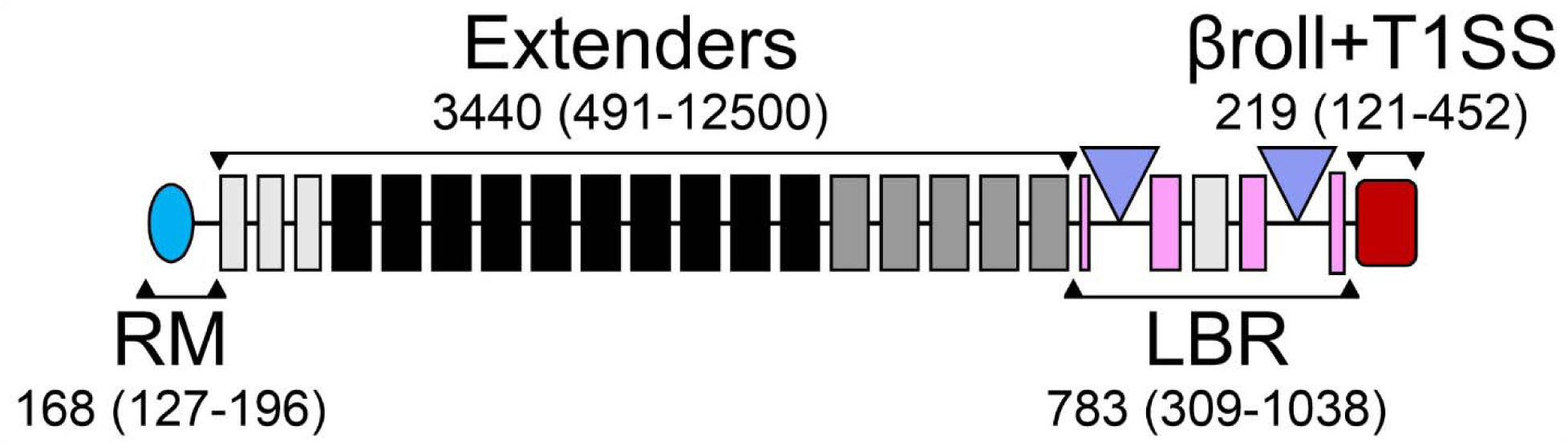
Schematic showing the general characteristics of RTX adhesins. The four sections common to most RTX adhesins are indicated with horizontal brackets and include the retention module (RM, cyan oval) at the N terminus, a string of extenders (black and grey rectangles), the ligand-binding region (LBR, with split domains (pink) from which ligand binding domains (blue triangles) protrude) and the C-terminal β-roll and type I secretion signal (βroll+T1SS, dark red rectangle). The numbers indicate the average length of these segments in residues from a sample of 50 adhesins, with the numbers in brackets indicating the range.

Structure-function studies on RTX adhesins show that most of their domains are Bacterial Immunoglobulin-like (BIg) beta-sandwich modules that extend the distal ligand-binding region (LBR) outward from the bacterial surface to help contact binding partners ^3, 29–31^. LBRs that have been examined in detail typically have two or three LBDs projected outward from the adhesin by ‘split’ domains ^32^. An example is provided by flagellar regulated hemagglutinin A (FrhA), a key RTX adhesin within the pandemic-causing *Vibrio cholerae* classical O strain. FrhA mediates both biofilm formation in the marine environment and attachment to intestinal cells during cholera infection ^7,13^. This relatively short adhesin has two LBDs. The first is a carbohydrate-binding module (CBM) that targets fucosylated surface glycans. The binding of this domain to erythrocytes is completely blocked by 5 mM fucose ^33^. The second is a peptide-binding domain (PBD) and sub-millimolar concentrations of the tripeptide Tyr-Thr-Asp-COOH were effective at blocking binding to both erythrocytes and epithelial cells ^7^. A second example is the LAP protein from *Aeromonas hydrophila*. This Gram-negative opportunistic pathogen is found in fresh and brackish waters ^34^ and one of its three putative LBDs is a CBM that binds to the fucosylated Lewis B and Y antigens ^8^. Again, millimolar concentrations of free fucose were found to block the binding of this CBM to a wide variety of targets such as yeasts, diatoms, erythrocytes, and human endothelial cells.

The examples above demonstrate that binding can be disrupted by natural ligands, but small molecule ligand mimics can be metabolized at slower rates, and they have already been demonstrated to be useful in blocking the initial colonization of bacterial targets, thereby preventing biofilm formation. For example, mannose-based antagonists that block FimH, an adhesin from *Escherichia coli*, can disrupt binding to mannose-containing glycoproteins in the uroepithelium, thereby reducing urinary tract infections ^35, 36^. This approach could be an alternative or adjuvant to antibiotics, which are rapidly losing their effectiveness ^37^. However, bacterial species can have more than one RTX adhesin. For example, *Pseudomonas fluorescens*, a soil bacterium that is rarely pathogenic ^38^, has two. Here, the long adhesin protein (LapA) and the medium adhesin protein (MapA) have complementary roles in adherence and the formation of biofilms ^20^. A second example is the pathogenic bacterium *Salmonella enterica*. The adhesin BapA is important for biofilm formation ^39^ whereas SiiE binds to glycans on epithelial cells ^12^.

These genes are located on chromosomal pathogenicity islands that are sometimes laterally transferred or not found in all strains of a species ^40^, which suggests that the complement of adhesins might vary by strain within a species. Therefore, to develop strategies to block colonization by a bacterial pathogen species, it is important to know the range of adhesins they make, how they are distributed between strains, and the variety of LBDs they collectively possess.

Due to their large size and repetitive nature, RTX and other large fibrillar adhesins are often poorly annotated in genome databases. Indeed, short-read assemblies cannot reliably be used for documenting long, tandemly repeated proteins ^41^. Here we have relied on the increasing availability of long-read genome assemblies to provide high-quality data with which to discover and characterize new adhesins, with a focus on RTX adhesins. Once large proteins are identified, InterProScan ^42^ is used to identify their domains using an integrated set of predictive models from multiple domain annotation databases. Although some key adhesin domains are frequently missed, BIg domains, as well as the N-terminal retention domain and the C-terminal T1SS that bracket full-length RTX adhesins are frequently found. This flags them for further analysis using AlphaFold3 where the structures of all the adhesin domains are predicted, which enormously helps with their identification. Moreover, the discovery of split domains ^32^ has facilitated the recognition of LBDs, although their function and ligands often remain unknown. The application of this bioinformatic process can reliably find and sort the RTX adhesins from a bacterial species that are annotated on long-read genomic sequences.

Here we have illustrated this methodology by applying it to seven bacterial pathogens of humans and other animals. The two *Vibrio* species examined, *V. parahaemolyticus* and *V. vulnificus*, are prevalent in seawater and cause more infections in developed countries than *V. cholerae*, primarily through the consumption of contaminated seafood such as oysters ^43^. The two *Aeromonas* species, *A. salmonicida* and *A. hydrophila* are found in fresh and brackish waters and cause massive losses in fish farms, with the former also being found in seawater and the latter causing wound infections, bacteremia and gastroenteritis in humans ^44^. *Legionella pneumophila*, a pathogen of single-celled organisms, can cause a severe pneumonia called Legionnaires’ disease if aspirated from contaminated water ^45^. The final two, *Bordetella parapertussis* and *Acinetobacter baumannii*, do not appear to have environmental reservoirs. The former is a causative agent of whooping cough that is restricted to humans ^46^ and the latter is causing increasing numbers of hospital-acquired infections ^47^. The acquisition of antibiotic resistance and ability of these species to form treatment-recalcitrant biofilms often leads to problematic infections and high mortality rates ^48^. Our bioinformatic process can rapidly identify the dominant adhesins and their LBDs, which could lead to the development of agents to block colonization and biofilm formation on host tissues. This bioinformatic process can also be easily adapted to catalogue other large proteins of interest, such as multifunctional autoprocessing repeats-in-toxin (MARTX) toxins ^49^ and non-ribosomal peptide synthetases (NRPS) ^50^.

## Methods

### Selection of quality genomes from which to extract RTX adhesins

A pipeline was written in Python that uses the NCBI Datasets v2 API ^51^ to find genomes for a user-specified species. The pipeline identifies genomes assembled using long-read technology, fetches all unique protein sequences within those genomes, clusters those proteins by shared identity, then produces a human-readable report. NCBI Datasets v2 was compiled as a Python library with OpenAPI. The first step of the pipeline was to fetch assemblies at the level of “complete genome” or “chromosome” (excluding “contig” and “scaffold”) for the given species from NCBI’s Genome database. Assemblies that failed completeness checks or were noted as contaminated were excluded, as were those that failed the average nucleotide identity (ANI) test for automatic species determination ^52^. In the case of redundant assemblies (GCF and GCA accession with the same ID), only the assembly from RefSeq (GCF) was retained. Next, only genomes obtained using long read sequencing technology were included by filtering the ‘sequencing technology’ portion of the record for any of the following keywords: ONT, Nanopore, Flongle, MinION, GridION, PromethION, PacBio, SRMT, RS II, or Sequel.

Each step of this pipeline is adjustable through a human-readable config file, which is set up ahead of each run. The source code is publicly available (<u>GitHub</u>).

### Sampling of filtered genome assemblies

In cases where more than 50 assemblies passed the filter criteria, the pipeline selected a sample of up to 50 representative assemblies. Sampling was stratified by the country of origin and collection date of the DNA, using a fixed random seed for reproducibility. To maximize diversity, the sampling method avoided duplicates from the same stratum (combinations of country and date) unless it was necessary to reach the target sample size. In such cases, remaining slots were filled by re-sampling the remaining genomes.

### Protein sequences and clustering

For each assembly in the sample, the pipeline retrieved its gene annotation package (a table of genes and their associated name, symbol, accession, and locus) from NCBI. Protein-coding sequences were generated by extracting all unique accession numbers, then fetching their corresponding amino acid sequences as a single FASTA file. To account for point mutations and other minor variations across samples, the pipeline clustered proteins by sequence identity using MMseqs2 ^53^. Clustering was performed based on the 800 C-terminal residues with a sequence identity threshold of 80%. For each cluster, the most frequently observed sequence was selected as the representative sequence. In the event of a tie, the longest sequence was chosen. The protein and assembly accessions within each cluster were presented alongside the representative sequence for downstream analysis.

### Identification of RTX adhesins within the output from the pipeline

Clusters containing proteins longer than 1,500 aa in length were selected for further analysis. Protein name annotations were used to exclude obvious non-adhesins. The representative sequence from candidate clusters was evaluated using the domain prediction tool InterProScan with default parameters. If domains commonly found in RTX adhesins were present, the sequence was selected for predictive modelling.

### Generation of models and verification of protein clusters

AlphaFold3 was used to model the structure of the representative protein from any protein cluster suspected of being RTX adhesin using the AlphaFold3 Server ^54^. If the sequence exceeded the 5,000-residue limit of the server, enough repetitive extender domains were removed to reduce the length below 5,000 aa. The model was visually inspected for the presence of the retention module and β-roll/T1SS domains. The boundaries of all domains and their structural similarity to known domain types was determined from the model and used to generate schematic diagrams of the adhesins. The sequence of the LBR plus the β-roll/T1SS, beginning with the first split domain and ending at the C terminus, was used in BLASTp searches ^55^ against all proteins in the report. Clusters were adjusted to include any additional sequences with > 75% identity to the LBR and to exclude those with <75% identity.

### Identification of LBDs by type

Domains were identified as vWFA domains by InterProScan. Putative CBMs and PBDs were identified by strand topology as described below. β-rolls were identified by their structure and the presence of at least one nonapeptide repeat motif (Gly-Gly-x-Gly-x-Asp-x-Φ-x), where x is any residue and Φ is a hydrophobic residue. Domains that did not resemble these three types were classified as unknown domains.

### Phylogenetics

The ‘Bacterial Genome Tree’ service ^56^, which uses RAxML version 8.2.12, at the Bacterial and Viral Bioinformatics Resource Center (BV-BRC) was used to generate a phylogenetic tree for the species mentioned in the manuscript. Representative chromosome-level assembly genomes were selected, and default parameters were used except that the number of genes selected for comparison was set to 500.

## Results

### Part 1: Semi-automated bioinformatics pipeline eases RTX-adhesin detection

#### Rationale behind using the pipeline to filter genome sequences

The difficulties encountered in identifying adhesins in sequence databases led us to develop a pipeline to simplify the process (Fig. 2). In total, over 55 adhesins were identified in the 29 Gram-negative bacterial species shown in Fig. 3, with the pipeline enabling rapid recognition and characterization of 35 RTX adhesins from seven species that are described in detail herein (Fig. 4). Briefly, the pipeline accepts a species as input, identifies genomes sequenced with long-read technology, retrieves all annotated protein sequences from those genomes, then clusters them based on sequence identity (Fig. 2). The steps are summarized in the Methods section, and the rationale behind each stage in the process is outlined below.

**Figure 2.**
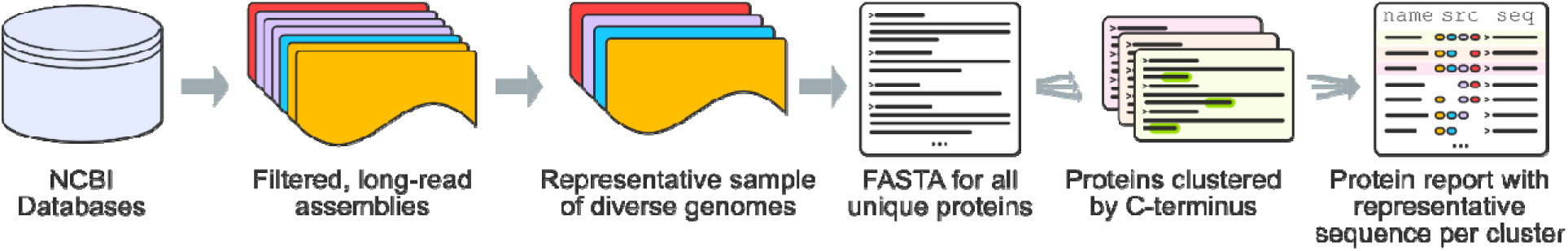
Flow of the bioinformatic pipeline used to identify unique RTX adhesin sequences for a given species. NCBI genomic databases are queried by species, and genome assemblies are then filtered for inclusion by sequencing technology, completeness, best average nucleotide identity (ANI) species match, and lack of notes for the assembly. A subset is taken from those filtered genomes to prevent over-sampling in cases where multiple assemblies are submitted from the same project, as well as to limit the size of the data for practical purposes. Clustering is performed on a specified stretch of residues with MMseqs2 to accommodate minor mutations across samples. Finally, a report is generated with per-cluster statistics, genome origin, and representative sequence indicating the most frequent isoform.

**Figure 3.**
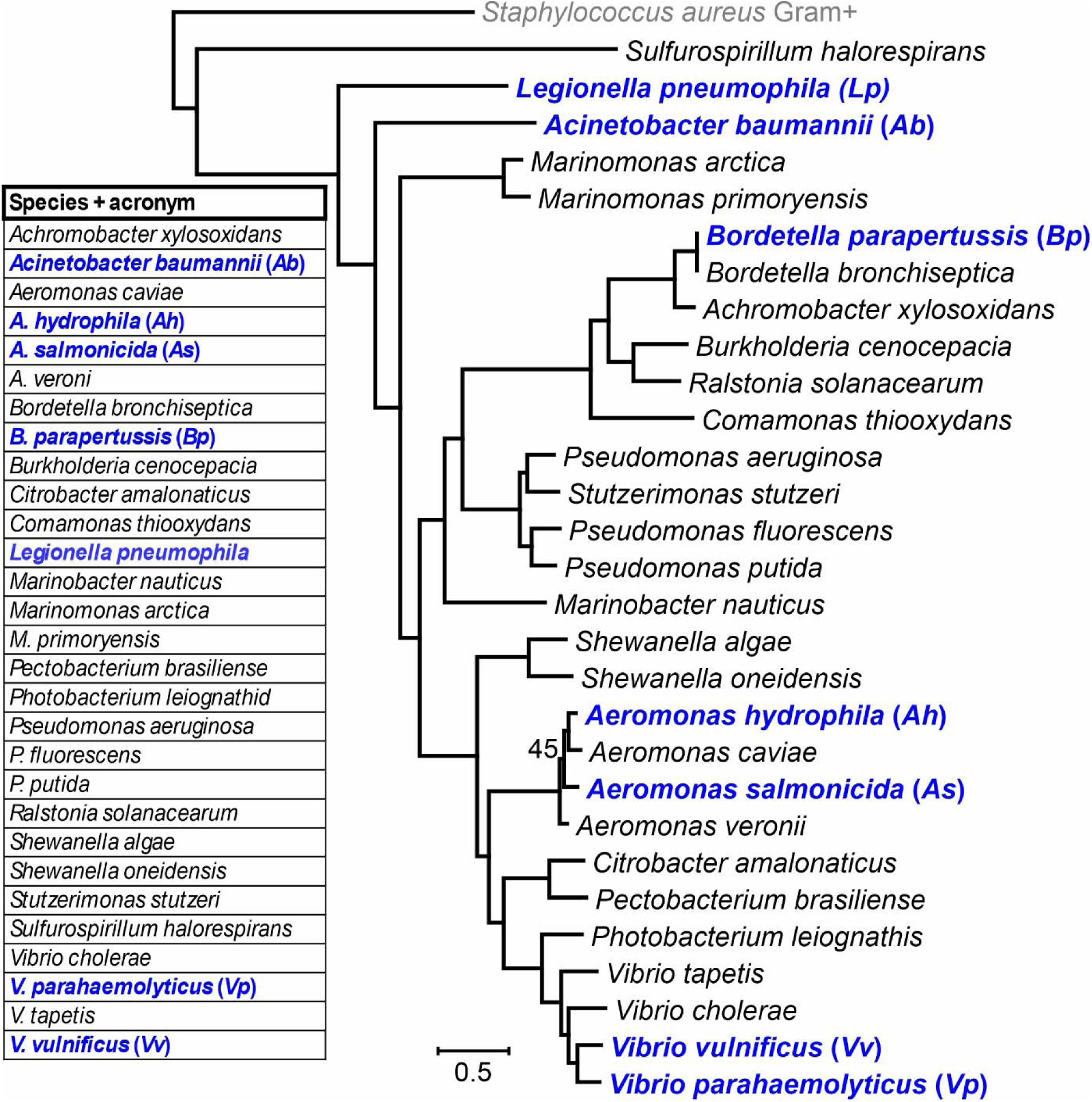
Gram-negative species (Kingdom Pseudomonadati) found to contain adhesins using the pipeline. Those whose adhesins are described in detail in this study indicated in bolded blue font. The species are indicated in alphabetic order in the table whereas the tree shows the phylogenetic relationships between the species, based upon 198 single copy genes (of 500 searched). Nodes did not change across all 100 bootstrap trials, excepting one within the *Aeromonas* grouping, so this is the only value shown. The tree was rooted using the Gram-positive species *Staphylococcus aureus*. The scale bar indicates a phylogenetic distance of 0.5 nucleotide substitutions per site.

**Figure 4.**
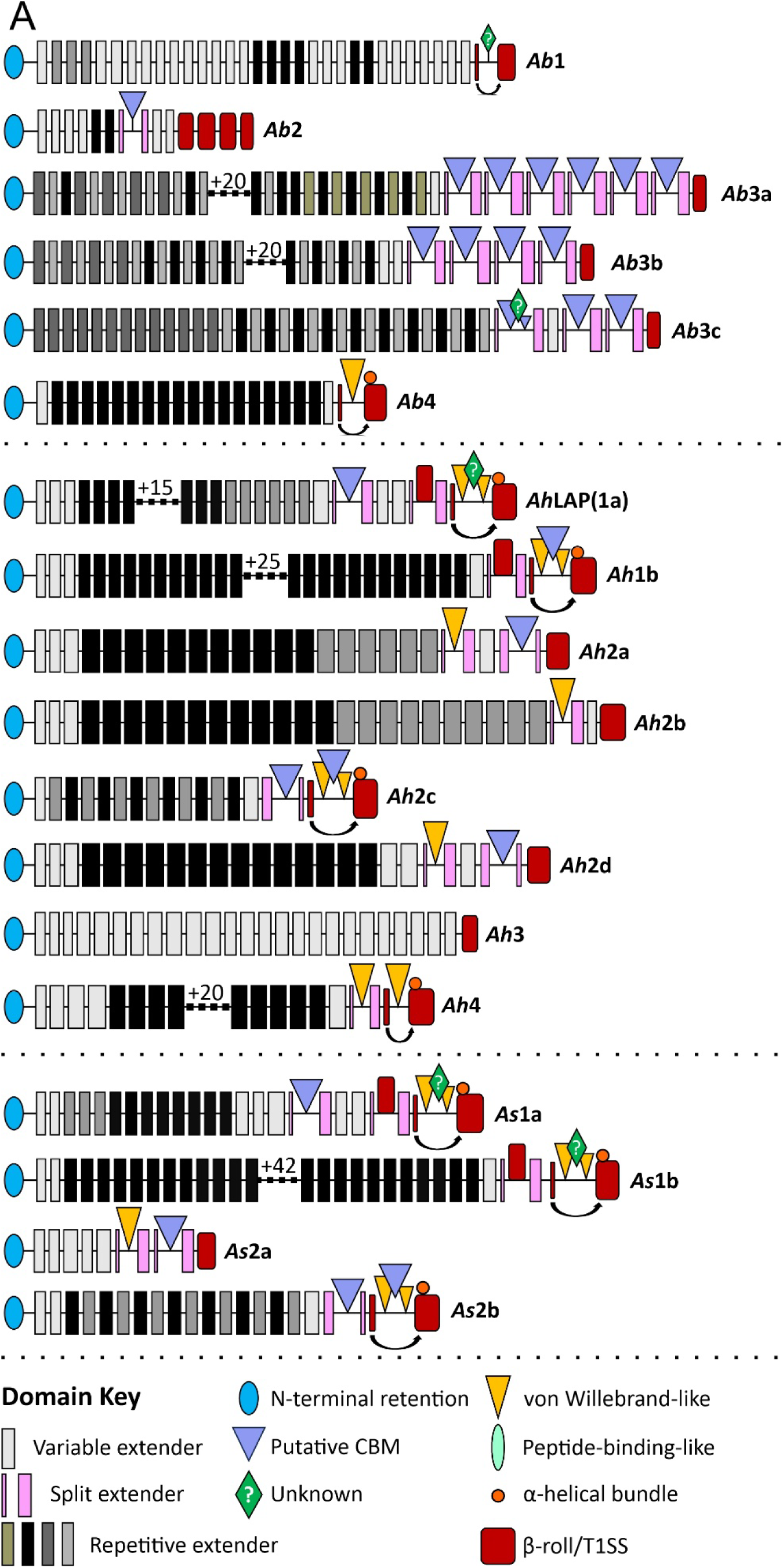
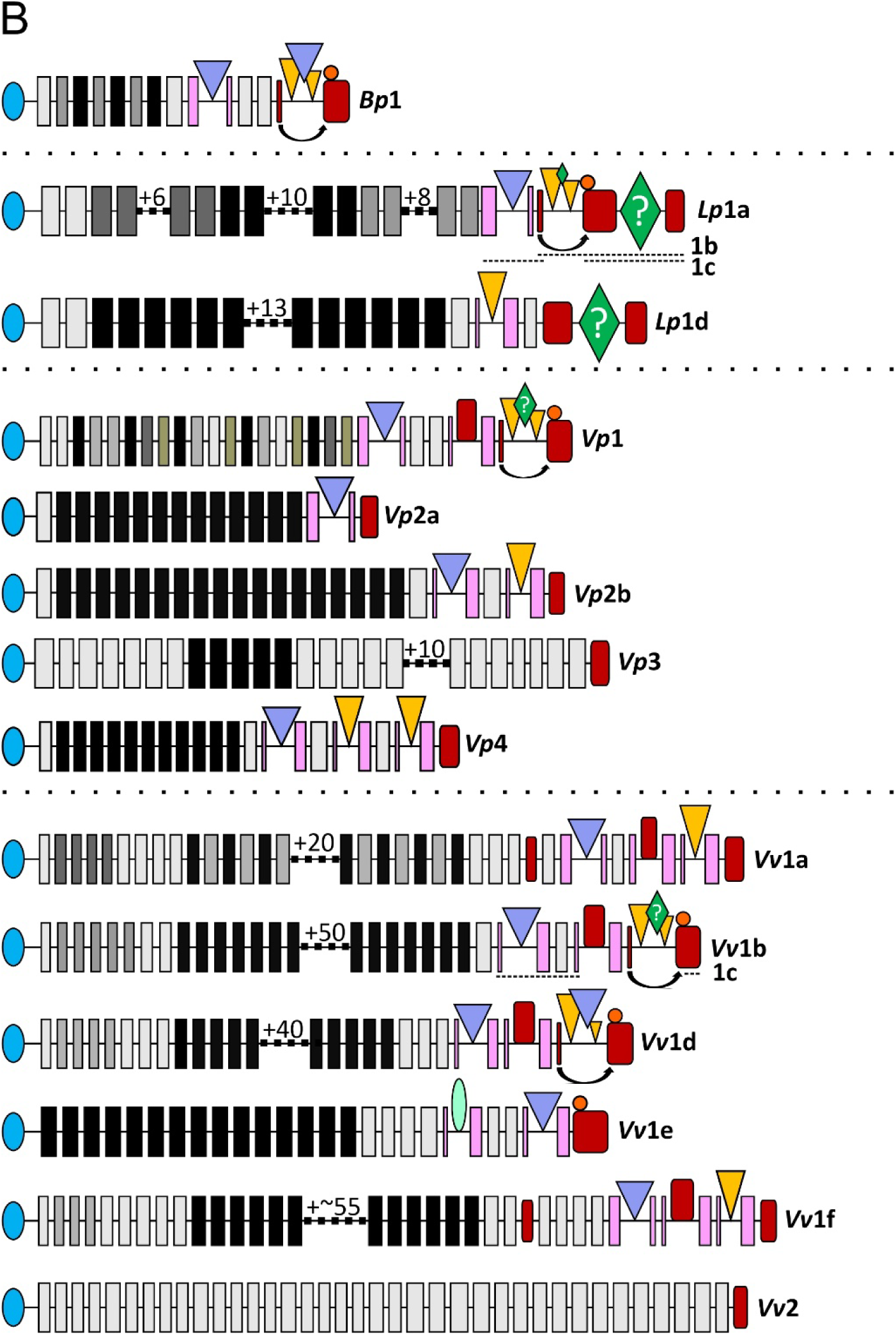
Domain maps for the adhesins found within the six bacterial species indicated in. Figure 1. Extenders are indicated with rectangles, where those in the lightest shade of grey are less than 80% identical to any other extender within the same protein. Darker shades of grey or black, as well as muted shades of green, are used to indicate groups of extenders that share at least 80% identity. A gap with a number indicates the number of extenders that have been omitted, which continue whatever pattern the extenders on either side show. The split extenders (pink) have a narrow bar at the left of the LBD if this projects after the first strand of the domain, or on the right if it projects just prior to the last two strands of the extender domain. The β-roll, vWFA and CBM domains can also act as split domains, indicated either by an arrow between the two segments (β-roll), or by two adjacent triangles with another projection (vWFA or CBM). The accession number of each representative sequence is given in Table 3. The dashed lines in *Lp*1a-c and *Vv*1b-c indicate portions of the LBR that are highly similar between the isoforms. The lengths of these isoforms differ (Table 3) due to variation in the number of extender domains, with the schematics representing *Lp*1a and *Vv*1b.

The first issue that came to light was that RTX adhesins are so poorly conserved between species that they cannot be reliably detected by BLASTp searches. For example, the highest scoring hit when the adhesin found in *Bordetella parapertussis* (*Bp*1) was compared to all adhesins in Fig. 4 was 29% identity to *Vp*2a over two thirds of its length. The same was sometimes observed even within a species, as the closest match to *Ab*3c is *Ab*3a, with only 35% identity. Despite this, they all share some common domains and are all far longer than the average protein. Therefore, survey methods that focus on these characteristics rather than sequence similarity are more likely to yield helpful results.

The second issue was that RTX adhesin ORFs were seldom assembled accurately due to their extreme length (approx. 4,500 to over 40,000 bp) and repetitiveness. Approximately a third of the adhesins shown in Fig. 4 contain more than 20 identical repeats, ranging in size from ∼250 to ∼650 bp per unit, that encode the tandemly arrayed BIg-like domains of the extender region (Fig. 4). Misassembly was particularly evident within genomes compiled from short-read sequencing (e.g. Illumina), while long-reads from single-molecule real-time (SMRT, Pacific Biosciences) or nanopore (Oxford Nanopore Technologies) sequencing usually spanned the entire gene and were properly assembled. Most submitted genomes are assembled from short reads, exemplified by *Acinetobacter baumannii* where only 833 of 40,037 genomes (2.1%) were high-quality assemblies from long reads (Table 1). The highly repetitive *A. baumannii* adhesin 3c (*Ab*3c) (Fig. 4A), provides a clear example of the problem: A BLASTp search of the GenBank non-redundant (nr) protein database identified 84 obvious homologues with >98% identity to the non-repetitive C-terminal region of the protein. Of these, only five were greater than 2,000 aa in length and all these originated from long-read assemblies. The longest (WP_269006110.1, 6,421 aa) has 46 identical 108-aa extender domains. Of the remaining 79 homologues, just over half (43) were described as partial, and the rest are unlikely to be complete or accurate. Therefore, to maximize the chances of finding properly assembled full-length RTX adhesins, the first step of the pipeline is to filter available genome assemblies to only include those that used long-read sequencing technology and meet quality criteria.

**Table 1.** Summary of bioinformatics analysis of selected bacterial species.

| Species acronym | *Ab* | *Ah* | *As* | *Bp* | *Lp* | *Vp* | *Vv* |
| --- | --- | --- | --- | --- | --- | --- | --- |
| **Genome Categories** |  |  |  |  |  |  |  |
| Sequenced genomes | 40,037 | 596 | 490 | 1,610 | 8,153 | 12,918 | 2,591 |
| Complete or chromosome<sup>1</sup> | 1,048 | 100 | 58 | 92 | 185 | 225 | 40 |
| Long read + filters<sup>2</sup> | 833 | 43 | 36 | 90 | 114 | 127 | 30 |
| Number sampled | 50 | 43 | 36 | 50 | 50 | 50 | 30 |
| **Protein Categories** |  |  |  |  |  |  |  |
| Total Unique Proteins | 34,220 | 84,343 | 46,459 | 4,652 | 32,171 | 83,972 | 60,743 |
| Clusters ≥ 800 aa | 358 | 567 | 409 | 118 | 264 | 579 | 463 |
| Clusters > 1500 aa | 66 | 64 | 39 | 13 | 32 | 67 | 57 |
| Unique RTX-adhesins<sup>3</sup> | 6 | 8 | 4 | 1 | 4 | 5 | 7 |

<sup>1</sup> Excludes assemblies that are at the contig or scaffold assembly level.
<sup>2</sup> Excludes genomes from species that did not meet the Average Nucleotide Identity criteria (ANI) of 96% identity over 90% of the genome <sup>52</sup>or that had warnings about completeness or accuracy. Includes genomes sequenced with either PacBio or Oxford Nanopore systems alone or in combination with short-read systems.
<sup>3</sup> Adhesins whose LBRs (from the start of the first split extender to the C terminus) have the same architecture.

If over 50 assemblies remain after quality filtering, they are subsampled by the pipeline for both practical and statistical reasons. Practically, protein sequences are fetched by assembly accession and processed locally. This means that for some organisms, like *Acinetobacter baumannii* which has 981 assemblies meeting our criteria, processing all of them would greatly inflate file size, run time, and server load to NCBI while yielding diminishing returns in analytical value. Additionally, large sequencing projects often submit many assemblies from a single strain or outbreak, causing their sequences to be overrepresented in the dataset. In *Vibrio parahaemolyticus*, 127 long-read assemblies passed our filter (Table 1) with twelve of these assemblies generated by a sequencing project investigating infection of a bivalve on the Pacific coast of Colombia ^57^. These data are excellent, but to consider all these assemblies would bias the analysis toward samples collected in a specific region that infect a specific organism.

Therefore, the subsample is generated using stratified random sampling of the filtered assemblies, drawing across country-and-date strata.

#### Clustering and filtering of proteins from the selected genomes

All annotated proteins encoded by the selected genomes are downloaded from the NCBI server by the pipeline. MMseqs2 then clustered all sequences at an 80% identity threshold based only on their 800 C-terminal residues. This C-terminal focus was adopted as even among isoforms with a highly similar ligand-binding region, their extender array often varies greatly in length or sequence. We had to consider a large enough window to compare more than the C-terminal secretory β-roll (120 to 450 aa) and after some experimentation, an 800-residue window was found to be sufficient. With all proteins clustered by their LBRs, we could then limit our search to sequences >1,500 residues, a minimal size for an RTX adhesin, thereby reducing the number of candidate sequences to screen (Table 1). The net effect was that rather than manually searching through hundreds of genome assemblies for adhesin-like sequences, a report for each species with fewer than 70 candidate sequences was quickly generated.

#### Identification of RTX adhesins from protein clusters and InterProScan

The prokaryotic genome annotation pipeline (PGAP) assigns algorithmically generated names for encoded proteins, which helps narrow our focus ^58^. These names are based on features detected in the sequence, so large well-characterized proteins such as ‘non-ribosomal peptide synthetase’ or ‘DEAD/DEAH box helicase’ can be further pruned from the dataset. For multidomain proteins such as RTX adhesins, however, the name is typically generated based on just one of its constituent domains. As such, adhesins are often assigned names like ‘Ig-like domain containing protein’ or ‘retention module-containing protein’, which is less reliable and necessitates further characterization.

The domain structure of any sequence suspected of being an RTX adhesin can be screened using InterProScan ^42^, and the web-based version will accept up to 100 sequences at once. A variety of domain types are recognized within each of the four regions of a typical RTX (Fig. 1) with the types that are commonly observed listed in Table 2. Domains within each region are sometimes not recognized, but at least two different domain types were found in the adhesins described herein (Fig. 4). Detailed examples of InterProScan results are provided in Supplementary Results and shown in Supplementary Fig. 1, with a complete list for all isoforms in Supplementary Table 1. This screening provides a rapid means of selecting proteins that have a domain structure consistent with RTX adhesins and excluding those that do not.

**Table 2.** Domains characteristic of RTX adhesins detected by InterProScan.

| Domain function | Automatic annotation name |
| --- | --- |
| Retention module | retention_LapA, BapA_N, |
| Extenders | Big_#, Ca_tandemer, Cadherin_#, CalX-beta, CalX-like, Ig-like_fold, LapA, tand_rpt_95, T1SS_rpt_143, VCBS_repeat, etc. |
| Ligand-binding | PA14 (rarely), VWF_A, vWFA, vWA-like, beta-Roll |
| β-roll | HemolysinCabind, beta-Roll, Serralysin-like_metalloprot_C |
| T1SS | T1SS_VCA0849, Peptidase_M10_C |

#### Modelling and verification of protein clusters using AlphaFold3

The representative sequence from any cluster suspected of being an adhesin was modelled using AlphaFold3 ^54^. Any candidate that exceeds 5,000 aa in length (the limit imposed by the AlphaFold server at the time of writing) was modelled with enough internal repetitive extenders removed to reduce the length below this limit. These models are undergoing processing in ModelArchive ^59^ and will be available prior to publication. In the meantime, the *.cif files can be obtained by request from the communicating author. Generally, the domains that were not detected using InterProScan were reliably modelled in AlphaFold3. Proteins that did not contain the canonical domains found in RTX adhesins, which are, at a minimum, the N-terminal retention module, a tandem array of Ig-like extenders, and a C-terminal β-roll with a T1SS signal, were excluded. The boundaries between individual domains were ascertained from these models and used to draw the domain maps in Fig. 4.

Within protein clusters, the number and sequence of extender domains was found to vary widely between homologues (described in more detail in Part 2), whereas the RM, LBR and β-roll/T1SS sections were usually highly conserved. However, as the LBRs of the different adhesins were often longer than the 800 aa length used to generate the protein clusters, proteins were sometimes misclassified. So, to verify that all cluster members are indeed homologous, the entire portion of the representative sequence C-terminal to the extender array (beginning with the first split domain) was used as a query in a BLASTp search against all members of the cluster, using the accession numbers provided in the spreadsheet from the pipeline. A second BLASTp was performed using the accession numbers of the representative sequences from all the other clusters, to ensure that all homologues were included. Any isoforms that matched with >75% identity across this entire region, with the same domain architecture, were considered homologues.

#### Determination of the prevalence of specific adhesins

By tracking the number of times a member of a cluster is found across sampled genome assemblies, the pipeline also gives a sense of the prevalence of RTX adhesins, as well as their isoforms, in a species. For example, for *Ab*3c mentioned above, the dearth of hits is surprising, given that there are almost 1,000 long-read assemblies and over 40,000 assemblies total for *A. baumannii*. This suggests that *Ab*3c is an uncommon adhesin in this species and indeed, this isoform was found only three times in the 50 long-read genomes examined (Table 3).

**Table 3.**
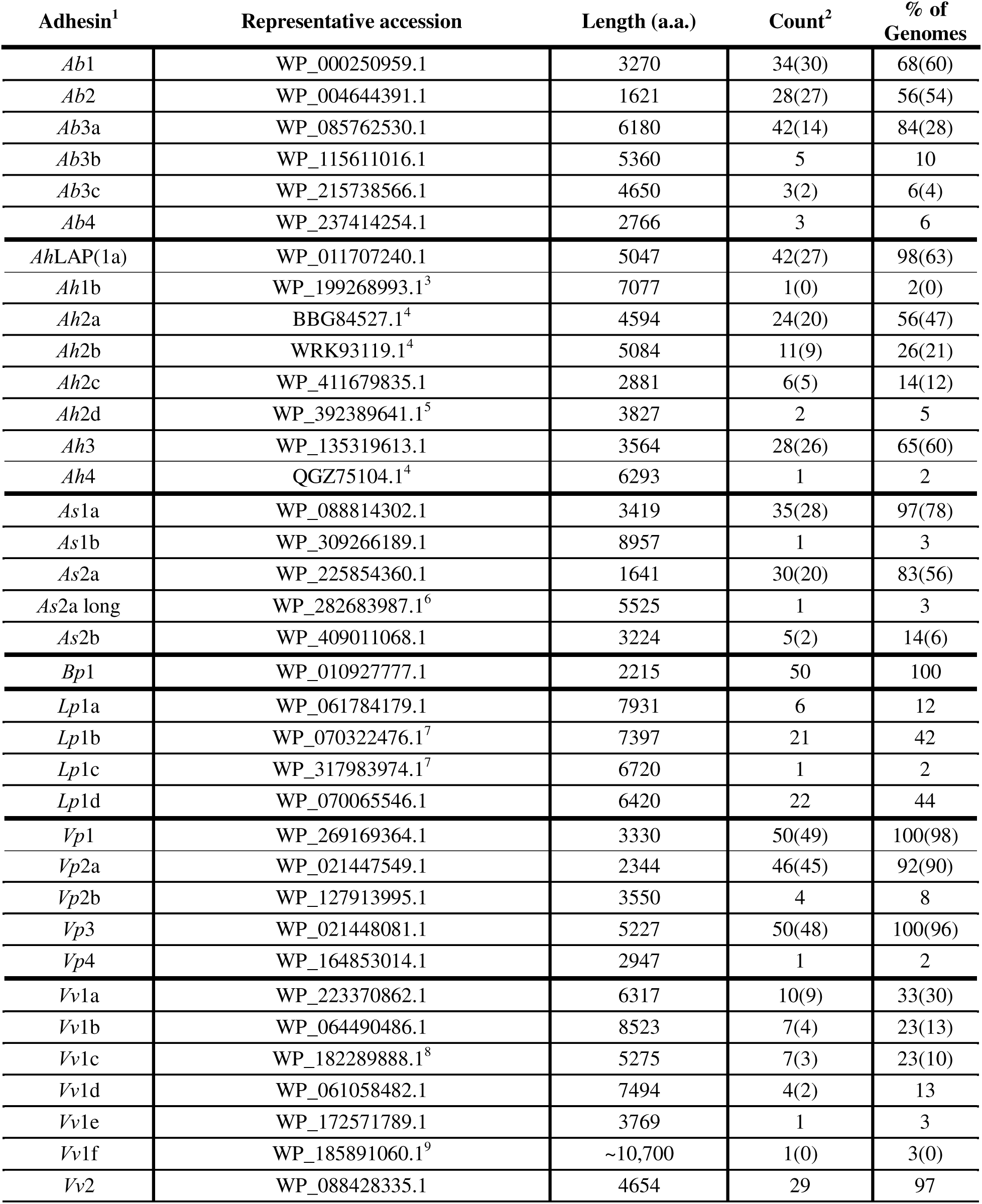
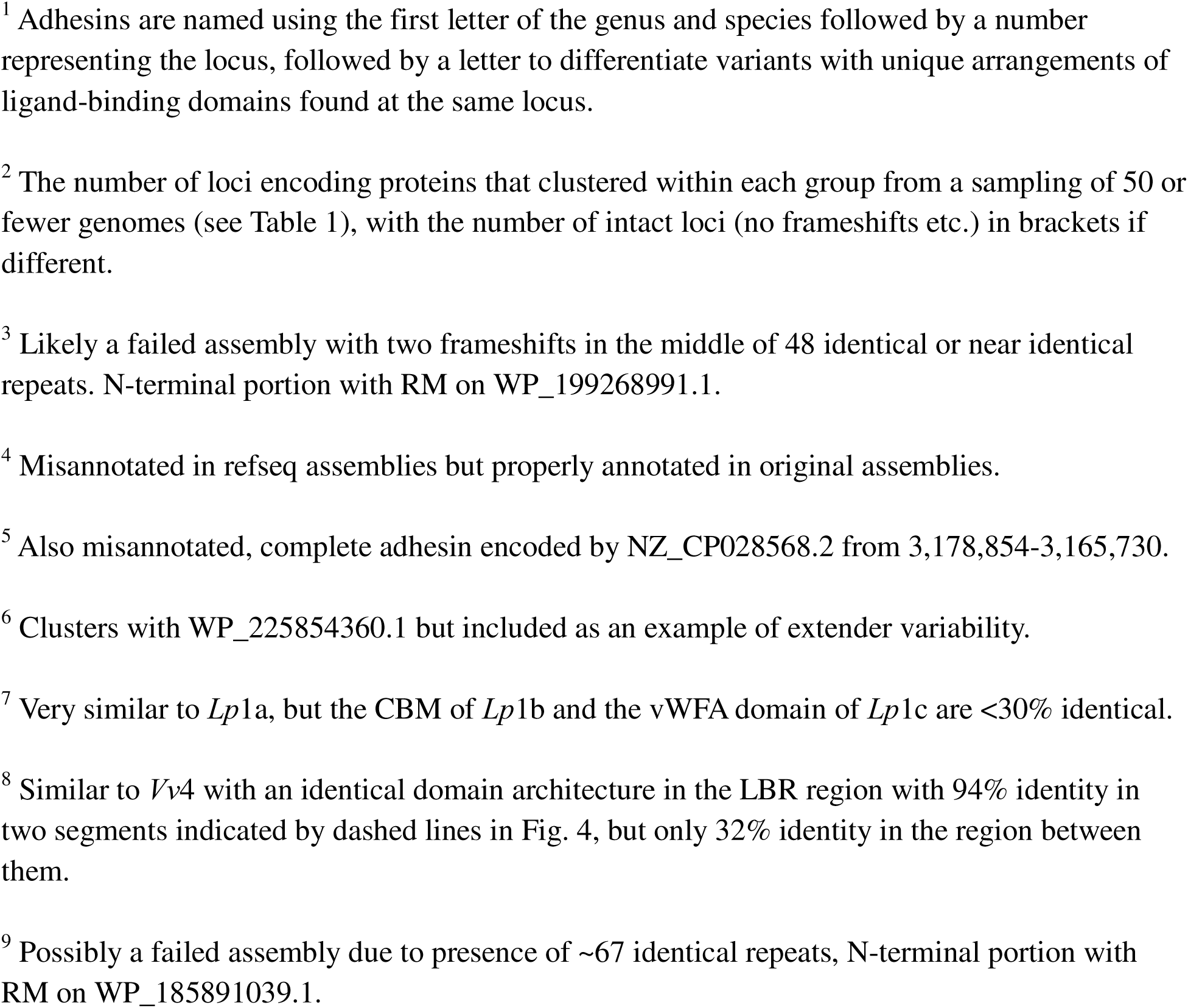
Accession numbers and parameters of representative proteins from each adhesin group.

| Adhesin<sup>1</sup> | Representative accession | Length (a.a.) | Count<sup>2</sup> | % of Genomes |
| --- | --- | --- | --- | --- |
| *Ab1* | WP_000250959.1 | 3270 | 34(30) | 68(60) |
| *Ab2* | WP_004644391.1 | 1621 | 28(27) | 56(54) |
| *Ab3a* | WP_085762530.1 | 6180 | 42(14) | 84(28) |
| *Ab3b* | WP_115611016.1 | 5360 | 5 | 10 |
| *Ab3c* | WP_215738566.1 | 4650 | 3(2) | 6(4) |
| *Ab4* | WP_237414254.1 | 2766 | 3 | 6 |
| *AhLAP(1a)* | WP_011707240.1 | 5047 | 42(27) | 98(63) |
| *Ah1b* | WP_199268993.1<sup>3</sup> | 7077 | 1(0) | 2(0) |
| *Ah2a* | BBG84527.1<sup>4</sup> | 4594 | 24(20) | 56(47) |
| *Ah2b* | WRK93119.1<sup>4</sup> | 5084 | 11(9) | 26(21) |
| *Ah2c* | WP_411679835.1 | 2881 | 6(5) | 14(12) |
| *Ah2d* | WP_392389641.1<sup>5</sup> | 3827 | 2 | 5 |
| *Ah3* | WP_135319613.1 | 3564 | 28(26) | 65(60) |
| *Ah4* | QGZ75104.1<sup>4</sup> | 6293 | 1 | 2 |
| *As1a* | WP_088814302.1 | 3419 | 35(28) | 97(78) |
| *As1b* | WP_309266189.1 | 8957 | 1 | 3 |
| *As2a* | WP_225854360.1 | 1641 | 30(20) | 83(56) |
| *As2a long* | WP_282683987.1<sup>6</sup> | 5525 | 1 | 3 |
| *As2b* | WP_409011068.1 | 3224 | 5(2) | 14(6) |
| *Bp1* | WP_010927777.1 | 2215 | 50 | 100 |
| *Lp1a* | WP_061784179.1 | 7931 | 6 | 12 |
| *Lp1b* | WP_070322476.1<sup>7</sup> | 7397 | 21 | 42 |
| *Lp1c* | WP_317983974.1<sup>7</sup> | 6720 | 1 | 2 |
| *Lp1d* | WP_070065546.1 | 6420 | 22 | 44 |
| *Vp1* | WP_269169364.1 | 3330 | 50(49) | 100(98) |
| *Vp2a* | WP_021447549.1 | 2344 | 46(45) | 92(90) |
| *Vp2b* | WP_127913995.1 | 3550 | 4 | 8 |
| *Vp3* | WP_021448081.1 | 5227 | 50(48) | 100(96) |
| *Vp4* | WP_164853014.1 | 2947 | 1 | 2 |
| *Vv1a* | WP_223370862.1 | 6317 | 10(9) | 33(30) |
| *Vv1b* | WP_064490486.1 | 8523 | 7(4) | 23(13) |
| *Vv1c* | WP_182289888.1<sup>8</sup> | 5275 | 7(3) | 23(10) |
| *Vv1d* | WP_061058482.1 | 7494 | 4(2) | 13 |
| *Vv1e* | WP_172571789.1 | 3769 | 1 | 3 |
| *Vv1f* | WP_185891060.1<sup>9</sup> | ~10,700 | 1(0) | 3(0) |
| *Vv2* | WP_088428335.1 | 4654 | 29 | 97 |

<sup>1</sup> Adhesins are named using the first letter of the genus and species followed by a number representing the locus, followed by a letter to differentiate variants with unique arrangements of ligand-binding domains found at the same locus.
<sup>2</sup> The number of loci encoding proteins that clustered within each group from a sampling of 50 or fewer genomes (see Table 1), with the number of intact loci (no frameshifts etc.) in brackets if different.
<sup>3</sup> Likely a failed assembly with two frameshifts in the middle of 48 identical or near identical repeats. N-terminal portion with RM on WP\_199268991.1.
<sup>4</sup> Misannotated in refseq assemblies but properly annotated in original assemblies.
<sup>5</sup> Also misannotated, complete adhesin encoded by NZ\_CP028568.2 from 3,178,854-3,165,730.
<sup>6</sup> Clusters with WP\_225854360.1 but included as an example of extender variability.
<sup>7</sup> Very similar to *Lp1a*, but the CBM of *Lp1b* and the vWFA domain of *Lp1c* are <30% identical.
<sup>8</sup> Similar to *Vv4* with an identical domain architecture in the LBR region with 94% identity in two segments indicated by dashed lines in Fig. 4, but only 32% identity in the region between them.
<sup>9</sup> Possibly a failed assembly due to presence of ~67 identical repeats, N-terminal portion with RM on WP\_185891039.1.

Conversely, *Ab*1 was found 34 times. It should be noted that the initial numbers may be underestimated, as even with long-read genomes the chance of an assembly error leading to a frameshift increases in proportion to the length and repetitiveness of the adhesin ^41^. However, if identical frameshift mutations are observed in multiple genomes, that would indicate that the adhesin has been lost in these strains, as described below.

#### Identification and verification of gene loci

The pipeline also outputs protein accessions paired with their corresponding genome assembly and chromosome accessions. The genetic neighbourhood of the genes encoding members of a cluster can be visualized across assemblies using a function built into the pipeline. This function orients and aligns a scaled view of chromosomal regions of the clustered genes including a user-specified number of their up- and downstream neighbours. Supplementary Fig. 2 shows this for *A. salmonicida* isoform 1a, found in 35 of the 50 genomes surveyed. The ORF is intact in 28 of these genomes, with the same three genes on each side. Anomalous genes can be examined using the graphics display at NCBI, which revealed that the two missing genes upstream of 4^th^ adhesin from the top are annotated as pseudogenes due to frameshift mutations. Of the seven ORFs that are incomplete, six contain putative frameshifts (indicated with purple stars) or a transposon insertion (indicated by a bar in light red). This function was used to generate the data for all isoforms, providing the counts in Table 1 and the details in Supplementary Table 2.

Comparing the list of searched genomes against those containing a particular isoform allows for rapid identification of missing genes. For example, a thorough analysis of the 43 *A. hydrophila* genomes has revealed that adhesins are ubiquitous at two loci (*Ah*1 and *Ah*2) but not at two others (Fig. 5A). The genes flanking each isoform are listed in Supplementary Table 3. In some cases, these adhesins were either misassembled, misannotated, or contain potential frameshifts (Supplementary Table 2). As the N-terminal RMs are usually well-conserved at each locus, a BLAST search using a complete sequence as the query and the protein accessions from the cluster as the subjects revealed those that do not start at the same position. Misannotations can be differentiated from frameshifts by performing a tBLASTn search against the chromosome accession to locate the “missing” DNA.

**Figure 5.**
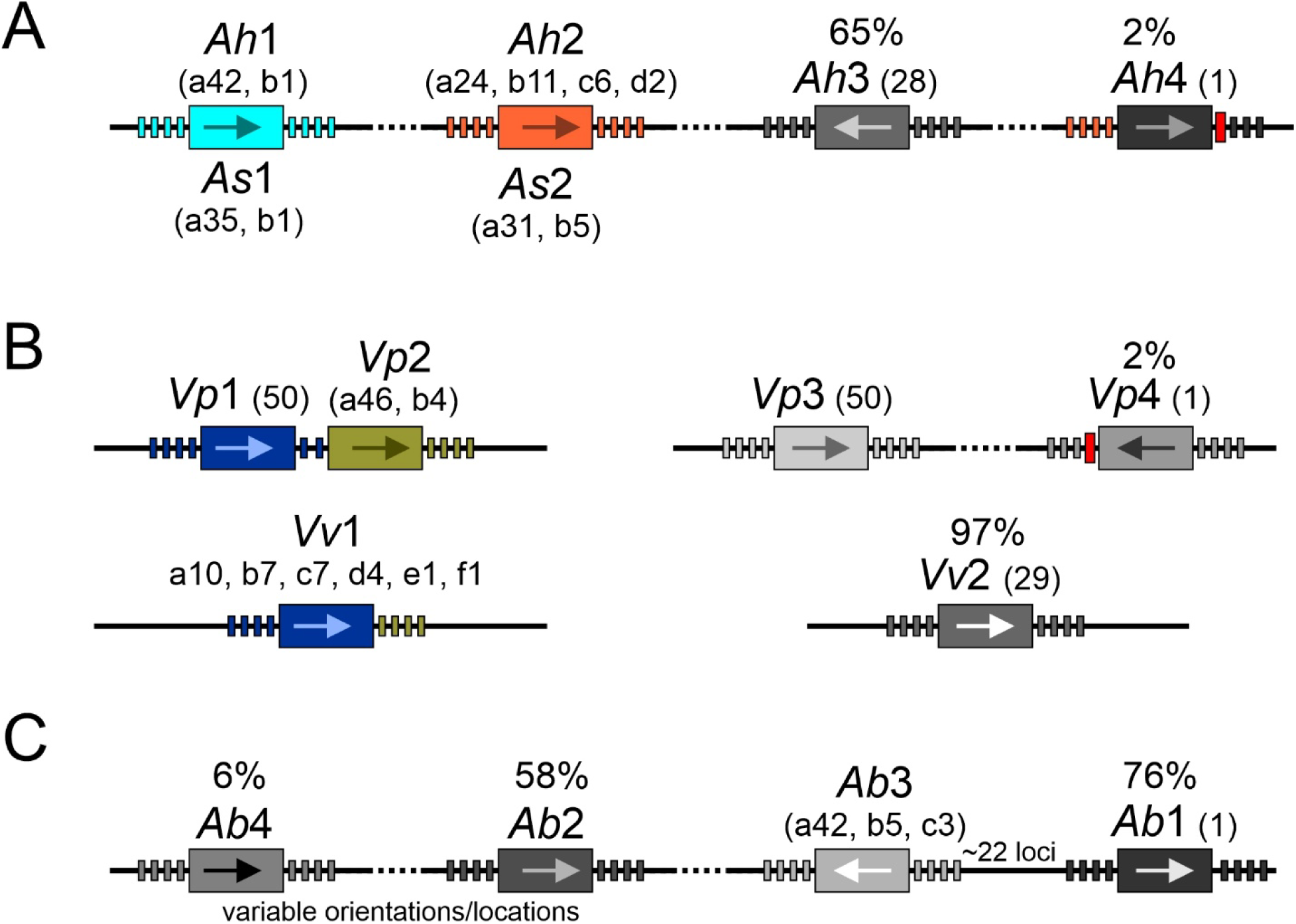
Schematic representations of adhesin loci from species with more than one locus (not to scale). A) Loci of the two aeromonad species B) Loci of the two Vibrio species found on chromosome 1 (left) and chromosome 2 (right). C) Loci of *A. baumannii*. Adhesins are represented with large rectanges and are labelled by locus and subtype as in Fig. 4, with an arrow indicating orientation. Dotted lines represent large distances between loci (>100 kb) and the four genes on either side are represented by small rectangles. Sequences derived from transposons are shown by slightly larger rectangles that are coloured red. Loci that have unique sets of flanking genes are coloured in various shades of grey. Those that share microsynteny are shown in the same colour. The names of the proteins encoded by these flanking genes are listed in Supplementary Table 3. The percentage of occupancy is indicated above each gene when it is less than 100%.

However, even when a frameshift is confirmed in the sequence via tBLASTn, distinguishing genuine biological frameshifts from sequencing or assembly artifacts remains a challenge.

Notably, most of the frameshifted sequences from *A. hydrophila* correspond to adhesins with large numbers of repetitive extenders. As well, all twenty-two of these frameshifts are unique (Supplementary Table 2), occurring at different locations within the gene (not shown). If the frameshifts were genuine biological features, they should occur at equal frequency in assemblies from both PacBio and Nanopore sequences. Instead, the ratio of frameshifts at the locus with the highest number of frameshifts (*Ah*1a) is higher in PacBio based assemblies (Supplementary Table 4). One explanation is that insufficient numbers of gene-spanning reads were obtained for accurate error correction using this technology. Therefore, it is difficult to ascertain whether these are genuine pseudogenes or simply genes with sequencing mistakes.

The abundant isoform at the second locus of *A. salmonica* (*As*2a) provides a contrasting example to that of the *A. hydrophila* loci. While the intact isoforms can be as long as 5525 aa and repetitive, most are like the short (1,641 aa) non-repetitive isoform shown in Fig. 4A, so misassemblies should be rare. However, frameshifts are found at ten of the thirty-one loci (Supplementary Table 2). Of these, nine share over 97% sequence identity with the locus encoding the short isoform with a frameshift at the same position in each (not shown). This is clear evidence that this frameshift is genuine and is not a result of misassembly. Analyses of this type are made easier due to the pairing of the proteins accessions with their corresponding genome accessions in the pipeline output.

### Part 2: General characteristics of the adhesins identified using the bioinformatics pipeline

#### Gene schematics reveal common themes and variations in adhesins

A total of 35 RTX-adhesins, possessing both an N-terminal RM and C-terminal β-roll/T1SS, were identified in the seven species. The schematic diagrams of these adhesins shown in Fig. 4 enable comparisons between proteins. The width of the extenders, split domain segments, and β-roll/T1SS segments are proportional to the lengths of these domains and illustrates how they vary in molecular weight. Numerous domain types serve as ‘split’ domains, which project putative LBDs away from the main axis of the adhesin, the most common of which are the BIg split extenders shown in pink. Regular extenders lie between the RM and LBR and their numbers vary widely, from five in *As*1 to over 70 in *Vv*6.

The cartoon model of one of the shortest (1,641 aa) adhesins identified, *As*2a, serves to illustrate the utility of the schematic diagrams (Fig. 6A). Here, both the vWFA (dark yellow) and CBM (purplish blue) can be seen to project out from the split domains. This adhesin also illustrates a frequent variation whereby proteins that have the same domain architecture and share >75% identity throughout the LBR can have very different extender regions. Here, the short isoform (*As*2a) has only five variable extenders, whereas the longest isoform (*As*2a long) has 37, and only the first two share any appreciable sequence identity. The opposite is sometime seen, as with *Ah*2a and *Ah*2d (Fig. 6B). The two adhesins share 97% sequence identity over most of their length and have the same domain architecture in the LBR, but the entire region from the last extender, through the entire LBR and partway into the β-roll was likely swapped in from another adhesin as the sequence identity drops to 28%. Examples of even smaller swaps are *Lp*1a, b and c (Fig. 4B and Table 3). These three highly similar proteins have the same domain architecture but the CBM of *Lp*1b shares only 28% identity with the other two, and the vWFA domain of *Lp*1c shares less than 30% identity with the other two.

**Figure 6.**
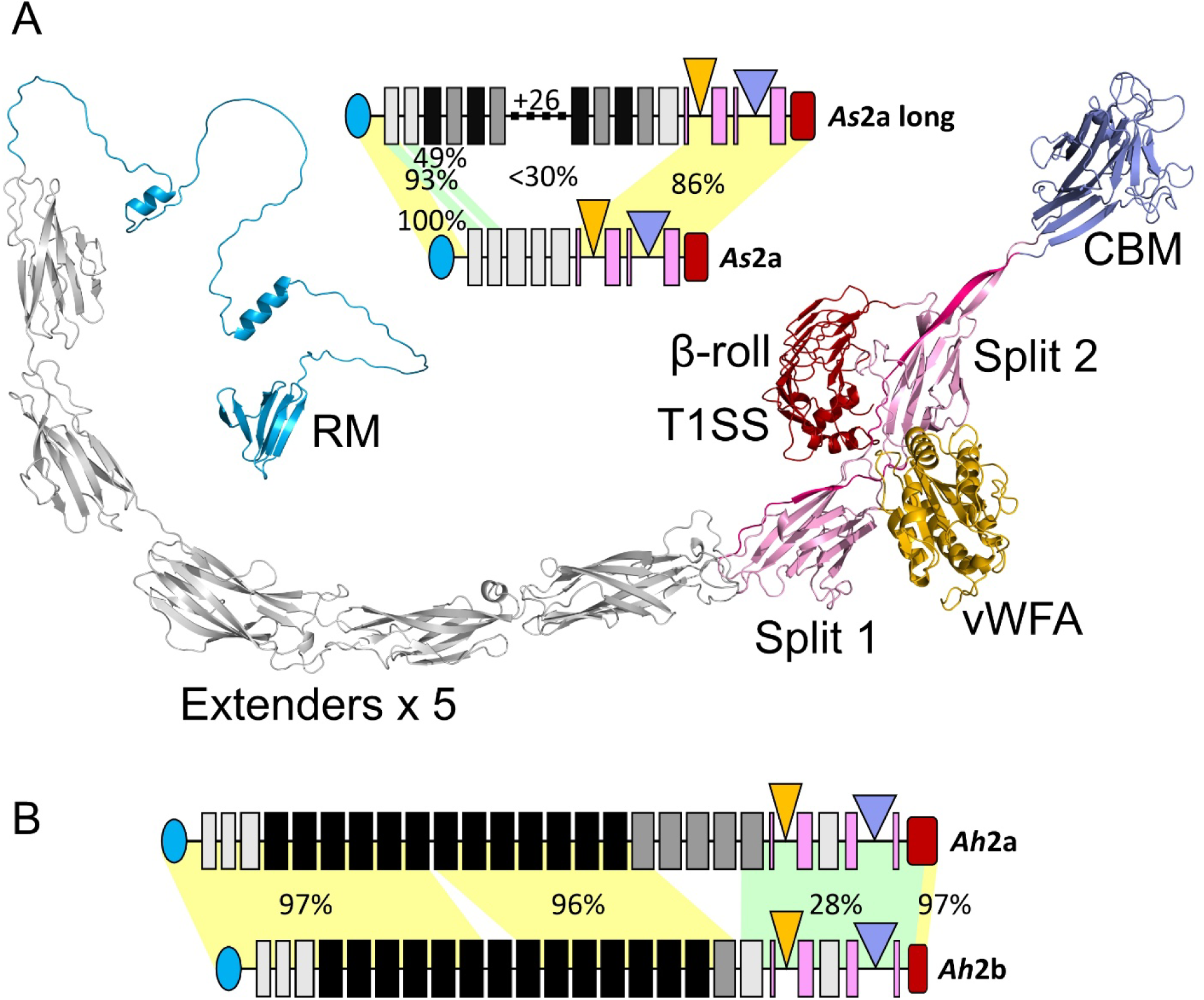
Comparison between structural and schematic depictions of adhesins and different patterns of sequence conservation. **A)** Cartoon depiction of the AlphaFold3 model of *As*2a with domains coloured as in Fig. 4, except that the first segment of the split domains are darker pink. The schematic of this short isoform is included, along with a schematic of the longest isoform of this type. The percent identity between sections of the two isoforms are indicated atop pale shading indicating the shared segments. **B)** Schematics of two *Ah* adhesins with percent identity indicated as in A.

#### Characteristics and variations of RMs and extenders

The RM domains of all adhesins described herein fall within a narrow length range of 127-196 aa (Fig. 1), with some variation in the number of strands within the RM and minor variation in the length of the linker. This domain was always followed by a region containing a string of extender domains, which can vary widely in number, with only five as in *As*2a to 73 as in *Vv*1f. The sequence similarity between extenders from different adhesins, and even between extenders within the same adhesin, is often so low (<25%) that they cannot be reliably aligned. Despite this, they all share an Ig-like fold consisting of a two-layer β-sandwich with an odd number of strands (most commonly seven or nine) such that the N and C termini lie at opposite ends of the domain, allowing the extenders to form a chain. The similarity of the fold is evident in two extenders at opposite extremes of a ∼2-fold length variation. The 85-aa extender 7 from *Ab*1 and the 178-aa extender 5 from *Ah*2a (Fig. 4) share a pattern consisting of seven segments, coloured in rainbow hues and numbered from N to C terminus (Fig. 7A, B). Around the circumference, the segment pattern is as follows; 1 (blue), 2 (cyan), 5 (yellow), 4 (light green), 3 (dark green), 6 (orange), and 7 (red). The main structural difference between the two are that the smaller extender domain has segments consisting of short continuous β-strands (Fig. 7A), whereas the segments in the larger extender (Fig. 7B) are sometimes split into two strands with an intervening loop or short α-helix. Additionally, the longer extender has two additional strands (dark grey). Split extender domains (Fig. 4, pink) within the LBR also share this fold (not shown).

**Figure 7.**
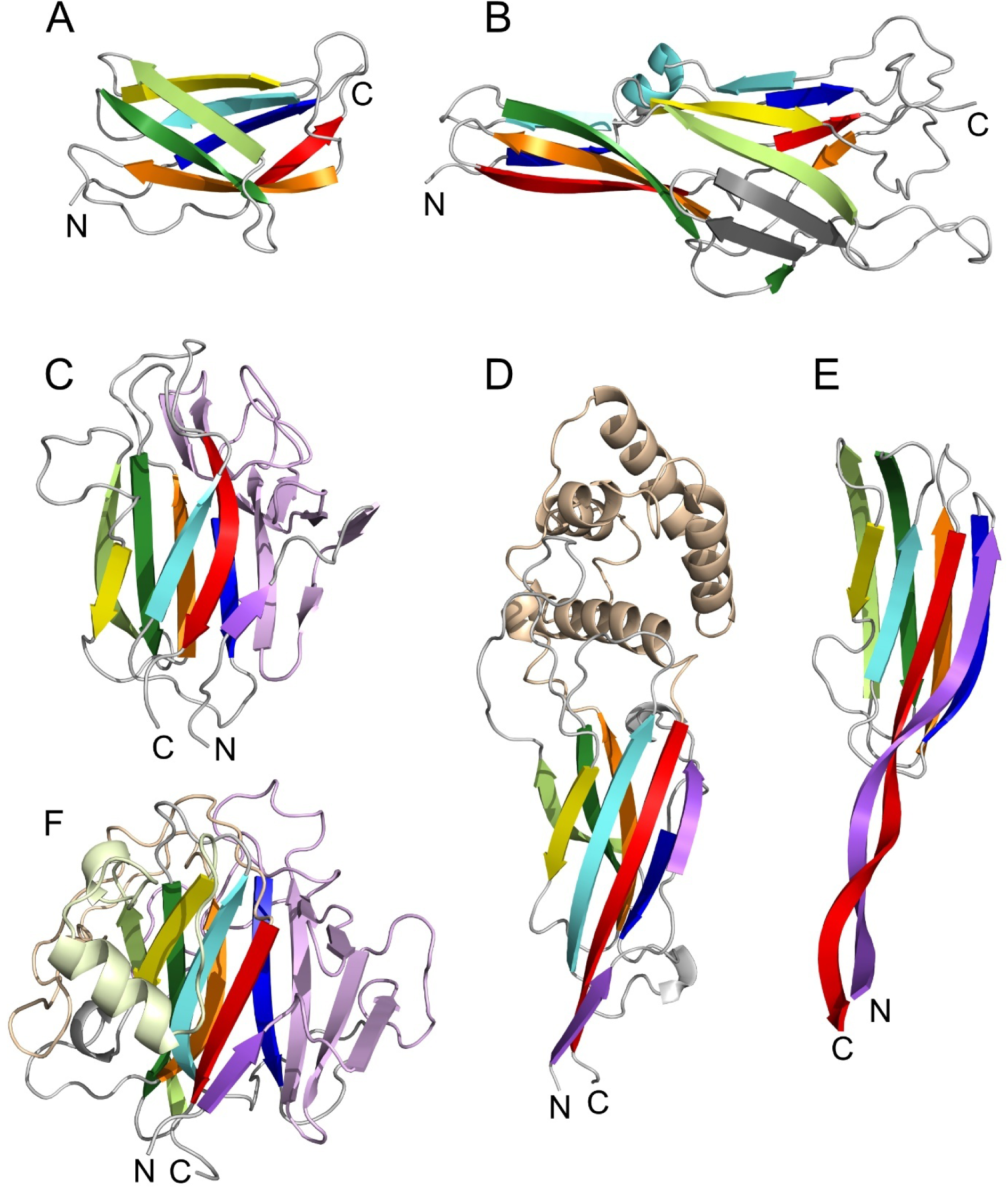
Shared features and differences between the models of representative domains identified as extenders and CBMs. A core set of seven strands are coloured in a rainbow pattern (blue, cyan, dark green, light green, yellow, orange, red) from N to C terminus for both domain types. The CBMs have eight strands, and the additional N-terminal strand is coloured violet. A) *Ab*1 extender 7 B) *Ah*2a extender 5 C) *Ah*Lap(1a) CBM D) *Ab*3c CBM-1 E) *Ab*2 CBM F) *Ah*2d CBM. Extra strands in the extender are coloured dark grey and in the CMBs they are coloured in a pale shade of the strand they follow, such as pale orange for the unknown helical domain in *Ab*3c.

The extenders that lie at either end of the extender region, along with those within the LBR, are usually dissimilar to other extenders in the same protein (Fig. 4, light grey), whereas those in the interior are often highly similar (Fig. 4, darker colours). The most common pattern is for adjacent extenders to be similar, as in *Vp*2b, where a set of 18 near-identical extenders is flanked on either side by a dissimilar extender. Another common pattern, found in *Ah*2c for example, is for two dissimilar extenders to form a pair that is repeated in series. More complex patterns occur, sometimes consisting of adjacent blocks of near-identical repeats exemplified by *Ah*2a, or consisting of a mixture of patterns as in *Ab*3c and *Vp*1. The other extreme is exemplified by *Vv*2, where each of the 37 extenders shares less than 80% identity with any of the others.

#### Characteristics and variations of ligand-binding regions

LBDs are generally recognized due to the presence of their companion split domains from which they project. The most common of these are the putative CBMs (Fig. 4, purplish blue triangles). These domains are highly variable, even within the same protein. For example, pairwise sequence identity between the six CBMs in *Ab*3a range from a low of 20% (CBM2 and CBM6) to a high of 56% (CBM1 and CBM3) (Supplementary Fig. 3). However, these domains can be recognized due to a common strand pattern (Fig. 7C-F). They have at least eight strands and around the circumference, the segment pattern is as follows: 1 (violet), 2 (blue), 7 (orange), 4 (dark green), 5 (light green), 6 (yellow), 3(cyan), and 8 (red). They are occasionally recognized as a PA14 domain by InterProScan, a CBM domain originally identified in the protective antigen of the anthrax toxin that is also found within yeast and other eukaryotes ^60^.

The vWFA domains are also common (yellow triangles, Fig. 4). They are easy to recognize as they consist of a six-stranded β-sheet with three α-helices on each face (Supplementary Fig. 4A) and they sometimes act as split domains themselves to other domains (Supplementary Fig. 4B). vWFA domains are readily identified by domain search tools such as InterProScan. The rarer PBD domain is more difficult to recognize and to differentiate from CBMs, until the strand pattern is examined. The PBD has two sets of strands in sequence in the pattern 6-7-8-9-10-5-4-3-2-1 (Supplementary Fig. 4C), in contrast to the less regular pattern for the CBM given above. β-roll domains, whether internal or adjacent to the C-terminal T1SS, are also readily identified by InterProScan, in part due to the prevalence of the nonapeptide repeat (Gly-Gly-x-Gly-x-Asp-x-Φ-x).

### Part 3: The Adhesins from seven Gram negative bacterial species identified using the bioinformatics pipeline

#### Acinetobacter baumannii adhesins are variable and complex

The adhesins of *A. baumannii* were characterized in a 2024 study where they were grouped into four types based upon their extender domains ^61^. Here, we have taken a different approach by grouping them based upon their C-terminal LBRs and their genomic neighbourhood. By this method, we have found six different adhesins populating four distinct loci, with three isoforms containing multiple CBMs at locus 3 (Fig. 4A, Fig. 5C). Both *Ab*3a and *Ab*3b were classified as the same type of adhesin in the previous study as some of their extenders are identical. However, their LBRs share less than 50% identity, with *Ab*3a having six putative CBMs while *Ab*3b has only four. The highest similarity between any two of these CBMs is 60% (CBM1 of *Ab*3a to CBM1 of *Ab*3b). A third type (*Ab*3c) has three CBMs, shares little sequence similarity with the others outside of the RM, and one of the CBMs acts as a split domain for a domain predicted to contain a bundle of α-helices (Fig. 7D). Given this, it is possible that these LBRs have different ligand specificities. Additionally, these adhesins have more putative CBMs than found in any of the other adhesins we have characterized (Fig. 4).

The three other loci encode proteins with other types of LBDs (Fig. 4A). Locus 1 has an unknown domain consisting of two long antiparallel strands that form a potential ligand-binding site at their tip together with another strand (cyan) and one α-helix (yellow) (Supplementary Fig. 4D). The protein from locus 2 has a unique LBR architecture with a simple CBM lacking any additional strands that is projected well away from the protein by a split domain in combination with two long β-strands (Fig. 7E). This is followed by four apparent β-roll/T1SS domains in parallel (Fig. 4A).The final protein (*Ab*4) has a single vWFA domain that buds off a split β-roll domain. This is a common architecture in RTX-adhesins (Fig. 4), but it is typically found in combination with other LBDs.

Adhesin sequences are found at locus 3 in all 50 assemblies, but 25 of these (*Ab*3a) share the same frameshift (Supplementary Table 2), a clear indication that this isoform is being lost. The occupancy at the other three loci ranges from a high of 76% (*Ab*1) to a low of 6% (*Ab*4). The pattern of gene loss/gain varies by locus. For *Ab*1, the eight flanking genes are present in all the surveyed genomes, whereas for *Ab*2, only the four downstream genes are present when the adhesin is absent, and for *Ab*4, the eight flanking genes are absent when the adhesin is absent. Together this suggests that adhesins are highly variable in some species and can be gained, lost, or altered at a high rate.

#### The single Bordetella parapertussis adhesin is highly conserved

An RTX adhesin has been reported in *B. parapertussis* as being involved in biofilm formation and attachment to respiratory epithelial cells ^62^. Here we show that this RTX adhesin is the only one in this species, found at a locus that does not share microsynteny with any of the loci from other species (not shown). It has several LBDs; a CBM-like domain projecting from a split extender domain and a vWFA domain projecting from a split β-roll domain that also possesses a small helical bundle (orange circle). This vWFA domain also acts as split domain for a second CBM-like domain. This LBR resembles those of *Ah*2c and *As*2b except with two extender domains between the CBM- and vWFA-split domain pairs (Fig. 4B). Despite this, *Bp*1 is only 33% identical to these, in the region from the start of the β-roll split domain to the C terminus.

The conservation of *Bp*1 contrasts sharply with those of most of the other species. Surprisingly, 49 of the 50 protein sequences are 100% identical. The only difference in the 50^th^ sequence is that it has lost one of the three repetitive extender pairs, depicted with alternating grey and black rectangles in Fig. 4B. In other species, identical sequences are the exception rather than the norm.

#### Legionella pneumophila adhesins reside at a single locus

Here we show that there are two main LBR arrangements in this species (*Lp*1a-c and *Lp*1d, Fig. 4B), occurring with similar frequency (56% vs 44%, Table 3). They reside at a single unique gene locus (not shown), so each strain has but one RTX-adhesin. Both forms have a vWFA domain, but in *Lp*1a-c it is set off by an internal RTX-β-roll split domain, whereas in *Lp*1d, it protrudes from a split extender domain. These two vWFA domains share less than 30% identity. The only CBM is found in *Lp*1a-c, and version found in *Lp*1b is only 28% identical to the others. Similarly, the vWFA domain of *Lp*1c shares less than 30% identity with the others, whereas the rest of the LBR of these isoforms match closely. The unknown domain (green, Fig. 4) is a 12-membered helical bundle that is not found in any other adhesin identified herein. It is >92% identical between all these isoforms.

#### The adhesins of Vibrio and Aeromonas species share many features despite limited sequence similarity

The adhesins of the two most closely related genera in this study, *Aeromonas* and *Vibrio* (Fig. 3) have LBRs dominated by CBM, vWFA and internal RTX-β-roll domains (Fig. 4). Many of these share common patterns, such as *Ah*Lap(1a), *As*1a, *Vp*1 and *Vv*1b where it is only the extenders that vary. Three of these species also share adhesins that lack recognizable LBRs (*Ah*3, *Vp*3 and *Vv*2), consisting entirely of extenders, most of which are variable, flanked by RM and β-roll/T1SS domains. Collectively, these genes reside at two to four different loci (Fig. 5), some of which are common to each species, and some of which are uncommon (Table 1). Most of these loci have one or more components of the type I secretion system, such as the outer membrane protein TolC, in their immediate vicinity (bolded in Supplementary Table 3).

We have found eight distinct RTX adhesins in *A. hydrophila* (Fig. 4A), residing at four loci (Fig. 5A), where the first in our list is the previously characterized *Ah*Lap ^8^. It has a CBM-split domain combination that is followed soon after by an RTX domain projected from a split extender domain and then a vWFA domain sporting a small unknown domain that emerges from the main RTX domain. The CBM of this adhesin has been shown by glycan-array analysis to bind Lewis B and Y antigens ^8^. This was the dominant isoform as it was found in 98% of the genomes examined, but as mentioned above, 36% of these have unique frameshifts that could be genuine or could have arisen from assembly errors (Supplementary Table 2). A rare allele at this locus (*Ah*1b, one of 43 genomes) is missing the CBM but its vWFA domain acts as a split domain for another CBM domain.

The second *A. hydrophila* locus hosts four alleles with various combinations of CBM and vWFA domains (Fig. 4A, Fig, 5A). *Ah*2a and *Ah*2d, appear similar, but the sequences of their LBRs share only 36% sequence identity. These genes were misannotated in most reference assemblies (Supplementary Table 2) so alternative accessions or annotations are provided (Table 3). They were predicted to begin at a non-canonical start codon, partway into the RM, downstream of a canonical in-frame start codon. The adhesin found at the third locus (*Ah*3) entirely lacks an LBR, with all 25 extender domains sharing less than 80% sequence identity and it is only found in 65% of the genomes. The fourth locus (*Ah*4) was only found in one genome and given that the LBR is 100% identical to an adhesin found in *Aeromonas allosaccharophila*, it is likely that it was recently acquired via lateral gene transfer from this species.

The four distinct RTX adhesins in *A. salmonicida* have identical or near identical LBR architectures to those of *A. hydrophila* (Fig. 4A). This is not surprising given that these two closely related species share two loci (Fig. 5A). Some of the sequences share also significant sequence similarity, as the LBR and RM regions of *As*1a and *Ah*Lap are ∼70% identical and *As*2b is over 80% identical to *Ah*2c over the length of the protein. Others do not, as *As*2a and *As*1b are quite dissimilar to any of the *A. hydrophila* adhesins. The two shared loci are common to both species while the extender-only locus is absent from *A. salmonicida* (Fig. 5A).

We identified five distinct adhesins in *V. parahaemolyticus* and seven in *V. vulnificus* (Fig. 4B). Both species have extender-only adhesins, *Vp*3 and *Vv*2 that are similar to *Ah*3, but sequence identity is below 30%. In all three species, these extender-only genes are found at different loci and do not share flanking genes (Fig. 5). *Vp*1, *Vv*1b and *Vv*1c resemble *Ah*Lap and *As*1a (Fig. 4), but here too, sequence similarity is low. The two genera do not share microsynteny at any adhesin locus, but the two *Vibrio* species do. *V. parahaemolyticus* has two adjacent loci, separated by two genes, with one isoform, *Vp*1 at the first locus and either *Vp*2a or *Vp*2b at the second. One locus, along with the two intervening genes, has been lost in *V. vulnificus*, and six different isoforms can be found here (Fig. 5B). *Vv*1b and *Vv*1c are another example where most of the LBR is conserved (Fig. 4B, dashed lines) with an intervening portion with the same domain types (no underline) that is quite distinct, sharing only 32% sequence identity.

Rare isoforms are also found in these two *Vibrio* species. *Vv*1e was only found once at locus 1 and it contains a completely different C-terminal LBR in which there is a putative peptide-binding domain ^63^ and a putative CBM separated by two extender domains. This is a similar pattern of LBDs to that seen in the FrhA RTX-adhesin of *V. cholerae*, with the LBRs sharing 87% sequence identity. This pattern is also seen in one RTX adhesin of *Aeromonas veronii* ^33^. Given the location of *Vv*1e (Fig. 5B), it may have originated from *V. cholerae* whereby a section of that gene was incorporated into *V. vulnificus* locus 1 via homologous recombination. The rare *Vp*4 isoform, found only once at a distinct locus, is likely an import from *V. diabolicus* (see WP_374091237.1) as the adhesins are 95% identical and of the same length.

The first three loci of *V. parahaemolyticus* and the two loci of *V. vulnificus* are fully occupied (Fig. 5B), except for one genome in which there is a large deletion that includes the *Vv*2 gene (not shown). There are a few genes that contain frameshifts, but as each one is unique, they could be genuine or they could be sequencing/assembly errors (Supplementary Table 2). Therefore, most *V. parahaemolyticus* strains have three adhesins, whereas *V. vulnificus* strains have only two. The extender-only adhesin within each species is highly conserved, whereas the adhesin(s) containing LBRs are more variable.

## Discussion

By only considering genomes sequenced with long-read technology, applying strict inclusion criteria to those assemblies, and employing a sampling strategy that prioritizes diversity, we have found full-length sequences for the common RTX adhesins in each bacterial species considered and identified diversity of their loci across the target species. Furthermore, we have identified and annotated the domains responsible for adhesion. Linking these adhesins to specific strains may help explain phenotypic differences and even permit the tailored design of strain-specific blocking strategies. Because LBDs are the most variable regions and are responsible for host attachment, they represent the primary targets for such interventions. Thus one might expect antibodies raised against the LBR to be specific to a particular species or strain.

### The diversity of pathogens studied reflects the ecological niche for RTX adhesins

The selection of these seven bacterial pathogens highlights the broad ecological distribution of RTX adhesins, spanning open water reservoirs, clinical settings, and human respiratory tracts. The *Aeromonas* (*A. hydrophila*, *A. salmonicida*) and *Vibrio* (*V. parahaemolyticus*, *V. vulnificus*) species represent pathogens that transition between aquatic environments (fresh, brackish, and coastal water) and diverse hosts. In these genera, RTX adhesins are shown to function in varying salinities and temperatures, as they facilitate attachment to both aquatic surfaces and host tissues ^64–66^. These proteins facilitate colonization, leading to significant morbidity in aquaculture^67–69^ and severe illness in humans via attachment to wounds or intestinal tissue ^70, 71^.

*Acinetobacter baumannii* and *Legionella pneumophila* represent opportunistic pathogens whose virulence repertoire, including RTX adhesins, allows for persistent colonization in human-engineered settings. *A. baumannii* is an increasingly prevalent cause of treatment-resistant hospital-acquired infections ^72, 73^ and these proteins likely underpin its ability to form robust biofilms ^48^. Similarly, *L pneumophila* thrives in warm freshwater systems, and uses its adhesins to colonize the human respiratory tract following transmission through aerosolized droplets ^74^. There, it causes Legionnaire’s disease, a particularly life-threatening form of pneumonia ^45^.

Unique in our list of species is *Bordetella parapertussis*, which lives exclusively in human respiratory systems. Infection causes whooping cough and ∼95% of *Bordetella spp.* infections in the United States are *parapertussis* ^75^, which may be because unlike other its relatives, it lacks an effective vaccine ^76^. This highlights the need to develop new antibacterial strategies and by targeting its RTX adhesin, it may be possible to prevent initial attachment and subsequent infection.

### Modular Evolution and the Combinatorial Architecture of RTX Proteins

The modularity of RTX adhesins makes their detection and classification a complicated task. A minimal RTX adhesin would comprise an N-terminal retention domain, a flexible linker that can span the T1SS channel, one or more tandemly repeated Ig-like domains, and a C-terminal β-roll/T1SS recognition domain. Through our analysis of this adhesin family, we have noted the degree to which each of these elements is prone to variation. Indeed, at the domain level, there is a “mix-and-match” architecture to the adhesins as though they have been assembled from a choice of retention domain variants, different types and quantities of Ig-like extenders, and a selection of LBDs budding off from different “split” extenders that are present in different arrangements and positions in the LBR (Fig. 4). Even the C-terminal T1SS recognition domain has variable length, exemplified by the very short (135 aa) version in *Ab1* and the much longer (317 aa) version in *Vv*5. This combinatorial quality might reflect DNA shuffling under selection pressure for the host bacteria to colonize different ecological and host-surface niches.

At a sequence level, we notice very low pairwise identity between like domains, not only across adhesins, but within adhesins. For example, in *As*2a, the tandem array of Ig-like extenders is a mix of at least four different versions, two of which are shown (Fig. 6A). Such poor conservation supports the hypothesis that these play the simple structural role of projecting the binding domains away from the bacteria and toward their target. If these domains fold correctly, it is largely unimportant which residues they are comprised of. In the LBR of *Ab*1, six putative CBMs are present and the highest match in a pairwise identity matrix was 56%. Here the lack of conservation might support a different hypothesis: that these CBMs mutate adaptively to bind different host targets.

### Genomic stability in Bordetella parapertussis and the challenge of divergence

The presence of a single RTX adhesin isoform in *B. parapertussis* is unique among the species considered. Initially, we thought this was indicative of an error in our pipeline, but closer inspection revealed that *B. parapertussis* exhibits astonishingly low genomic diversity throughout the entire species. Fifty sampled *Bp* genomes contained an average of 4,221 protein-coding genes yet yielded only 4,652 unique protein sequences. By comparison, *V. parahaemolyticus* with an average of 4,756 protein-coding genes per genome had 83,972 unique sequences across fifty genomes (Table 1). This makes it likely that an inhibitor against the solitary RTX adhesin would be effective against all cases of this human pathogen.

The diversity of proteins in *Vp* is typical of the species considered. This diversity highlights a key challenge in sequence-based protein annotation: standard tools may fail to recognize domains that have undergone extensive divergence. Indeed, attempts to use InterProScan produced mixed results, occasionally missing retention domains and extenders, and almost entirely overlooking the putative CBMs. Custom-trained HMMs, structure-based methods applied to predicted models, or some combination thereof could be developed to better annotate these features. Improved detection will be an important first step toward functional studies, which are needed to understand to what degree the sequence variation in binding domains affects their affinity for ligands. More reliable identification of adhesins and annotation of their binding domains will therefore be essential in developing blocking strategies.

### Evidence of gene duplication and interspecies DNA uptake and incorporation

There is considerable plasticity in the complement and structure of RTX adhesins present in these bacterial species, consistent with horizontal gene transfer. Bacteria are adept at incorporating DNA that provides them with a selective advantage, which in this case would presumably be for the colonization and/or infection of various hosts. As the RTX adhesins were described in the Results section it was convenient to draw parallels between similar LBRs in different species and strains. For example, the combination of a peptide-binding domain and a CBM separated by two extender domains seen in *Vv*6 is reproduced in *V. cholerae* and *A. veronii*^32^. The domain arrangement in the LBR of *Ah*Lap where vWFA, RTX and putative CBM domains are present in that order is also seen in the LBR of *Vp*2 and closely resembles that present in four different *Vv* RTX adhesins (*Vv*1, *Vv*3, *Vv*4, and *Vv*5). A third example might be the tandemly arrayed putative CBM-split domain modules seen in some *Ab* RTX adhesins (*Ab*1, *Ab*2, and *Ab*3) that are similarly arranged in a *Citrobacter amalonaticus* RTX.

### RTX adhesins without an LBR

Three of the seven bacterial species (*Ah*, *Vp*, *Vv*) featured in Fig. 4 contain single RTX adhesins (*Ah*3, *Vp*3, *Vv*2) that completely lack LBDs. A fourth species (*Ab*) has a small insertion budding from the C-terminal RTX domain of adhesin *Ab*5 but is otherwise devoid of any LBR elements. These LBR-lacking RTX adhesins are not rare but are well represented in the genomes of these species and are typically paired with an LBR-containing RTX adhesin in the same genome (Table 3). For example, *Vv*2 is found in the same strains as *Vv*1. This situation is reminiscent of RTX adhesins in the classic strain of *V. cholerae* where FrhA is the RTX adhesin with a LBR containing both a peptide-binding domain and a CBM separated by two extender domains, and CraA is an RTX adhesin that completely lacks LBDs. The latter has been postulated to have a role in biofilm formation ^11^.

### Generalized use of the bioinformatic pipeline

The bioinformatic pipeline used here can be readily adapted to identify and characterize diverse protein families including large ones like MARTX toxins ^22, 77, 78^ and NRPS ^79, 80^. Indeed, these two protein types are frequently seen in the outputs alongside RTX adhesins in the >1,500 aa size category and feature a similar mix-and-match architecture. To facilitate broader use, we have simplified the installation and user configuration. The ability to pick out long-read genome assemblies makes it ideal for finding proteins with repetitive sequences. The parameters can be adjusted to cluster any section of the sequence, so any protein of interest could be evaluated to determine both its prevalence and degree of conservation within a species.

## Abbreviations

BIg: Bacterial Immunoglobulin-like
CBM: Carbohydrate-Binding Module
FA: Fibrillar Adhesins
FrhA: Flagellar-regulated hemagglutinin A
LapA: Long adhesin protein A
LBR: Ligand-Binding Region
LBD: Ligand-Binding Domain
MapA: Medium adhesin protein A
MARTX: Multifunctional Autoprocessing Repeats-in-ToXin
NRPS: Non-Ribosomal Peptide Synthetases
PBD: Peptide-Binding Domain
RM: Retention Module
RTX: Repeat-in-ToXin
T1SS: Type I Secretion System
vWFA: von Willebrand Factor A-like domain

## Acknowledgements

This work was funded by GlycoNet Collaborative Team Grant C-11 to PLD and SG, and NSERC Discovery Grant RGPIN-2022-03845 to PLD.

## Supplementary Results

The domain structure of candidate sequences can be ascertained by using InterProScan ^42^, accessed via the European Molecular Biology Laboratory - European Bioinformatics Institute (EMBL–EBI) website. In some instances, domains corresponding to all four sections of an RTX adhesin (Fig. 1A) are detected (Supplementary Fig. 1, *Ah*Lap(2a)). Here, the first section was identified as a retention module (RM) and Table 2 shows which feature names correspond to this and the other domain types. Nearly all the extender domains (Ex) in section 2 were detected, as was the β-roll and von Willebrand factor A-like domain (vWFA) domain within section 3. Both the β-roll and the T1SS signal of section 4 that mediate secretion were detected. The domains that were not detected were the CBM (usually missed but occasionally detected as a PA14 domain) and one of the extenders. Split domains are seldom detected. In a second example, *As*2a long, domain detection was very similar except the bulk of the extenders were not detected.

However, three were identified, two near the N terminus and one within the LBR. At least one extender was identified by InterProScan in all adhesins examined (not shown). In the third example (*Vp*2a), domains were not detected in region 1 or 4. However, there are gaps in the domain map, indicating the possibility of undetected domains. Indeed, when modelled, *Vp*2a clearly possesses both a retention module and a β-roll/T1SS. Therefore, any proteins with domain maps that resemble the three shown here were selected for modelling.

---

**Supplementary Figure 1.**
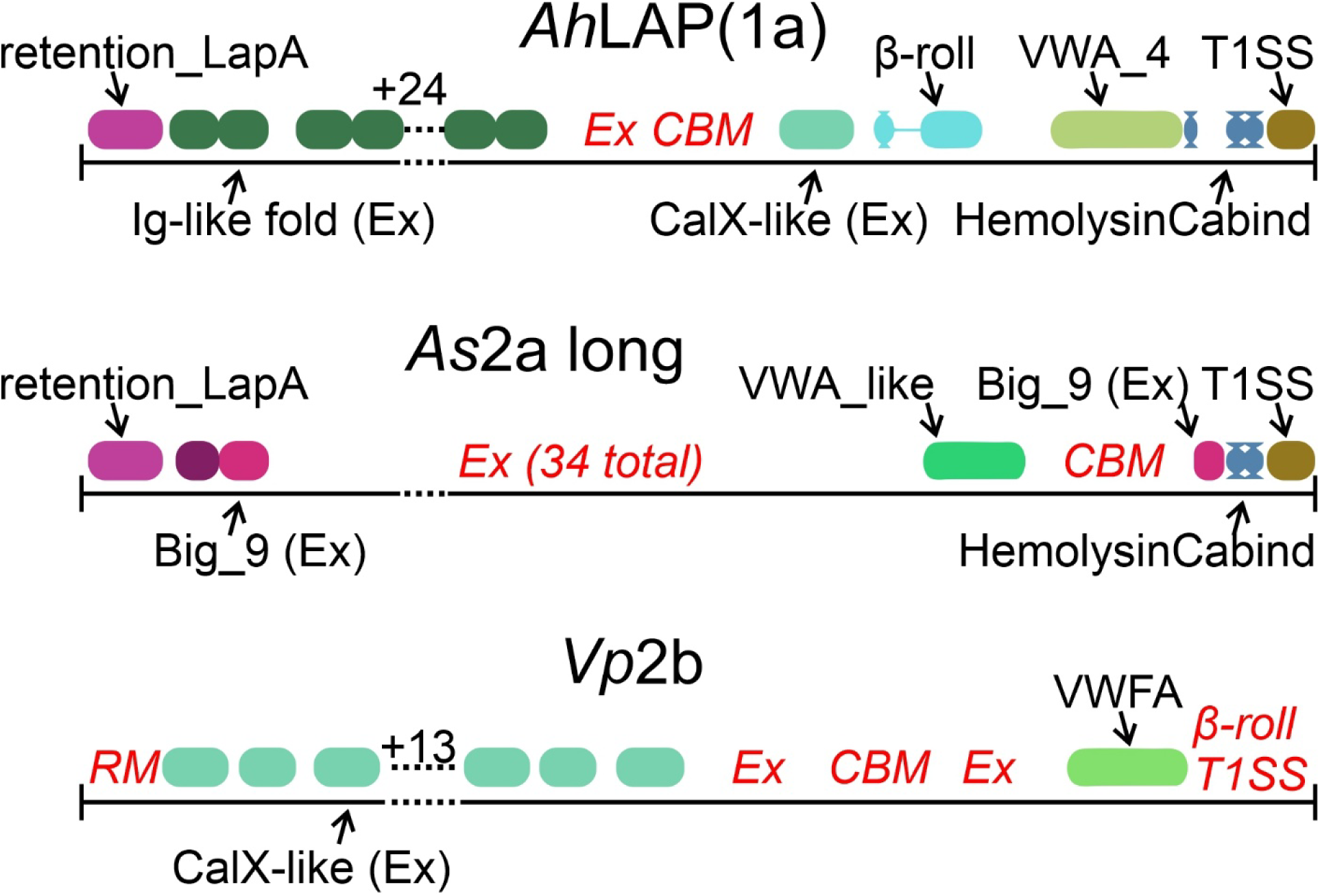
Representative InterProScan results for three RTX adhesins. The diagram was adapted from the actual output, including matching the colour, shape, and short descriptions of the domains found, but it has been simplified and modified. Domains that were not detected are indicated by red text in italics. Commonly detected domains are also listed in Table 2.

**Supplementary Figure 2.**
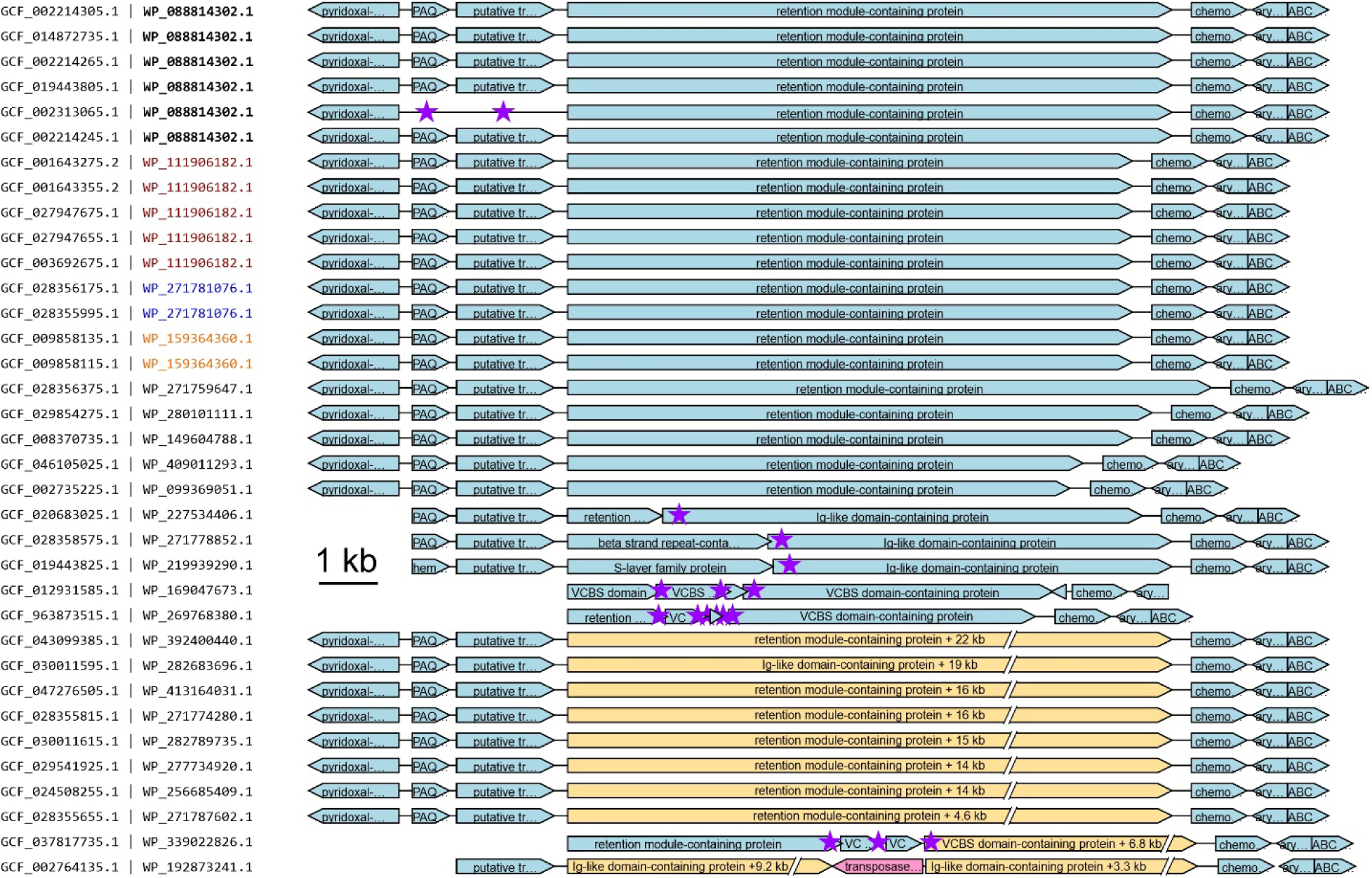
Schematics of the 35 *A. salmonicida* cluster members encoding isoform 1a (*As*1a) and their genomic neighbourhoods. This was output using a built-in function from the pipeline, which left-aligns the target proteins. It also shows a user-specified number of flanking proteins (three on each side here) as they appear within the genome in a scale diagram. The width of the bars is proportional to the length of each open-reading frame, except for the bars in yellow. The top twenty schematics were not adjusted, the next five were shifted to the right as the retention module was encoded separately, and the bottom ten were redrawn for clarity as these isoforms were much longer, so they are shown with a yellow broken line with the missing length indicated. Purple stars were added to indicate the locations of frameshifts and the transposon insertion was recoloured pink. The unique assembly accessions are paired with protein accessions, with the exemplar protein accession (identical protein in six genomes) in bold, and other identical proteins in dark red, blue, or orange font.

**Supplementary Figure 3.**
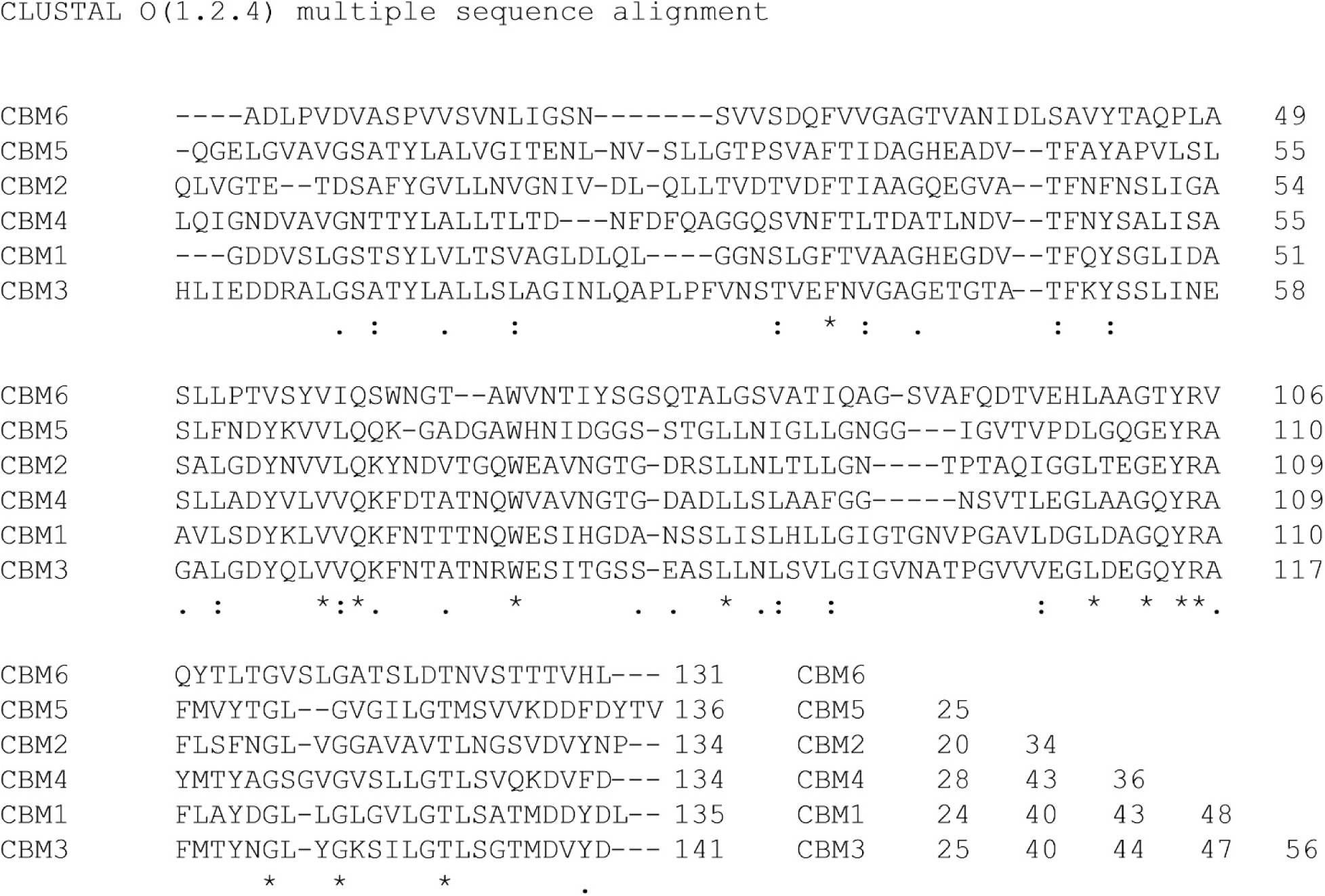
Alignment and percent identity matrix of the six CBMs of *Ab*3a. Asterisks indicate residues shared in all isoforms, with colons and dots representing conserved and semi-conserved residues, respectively.

**Supplementary Figure 4.**
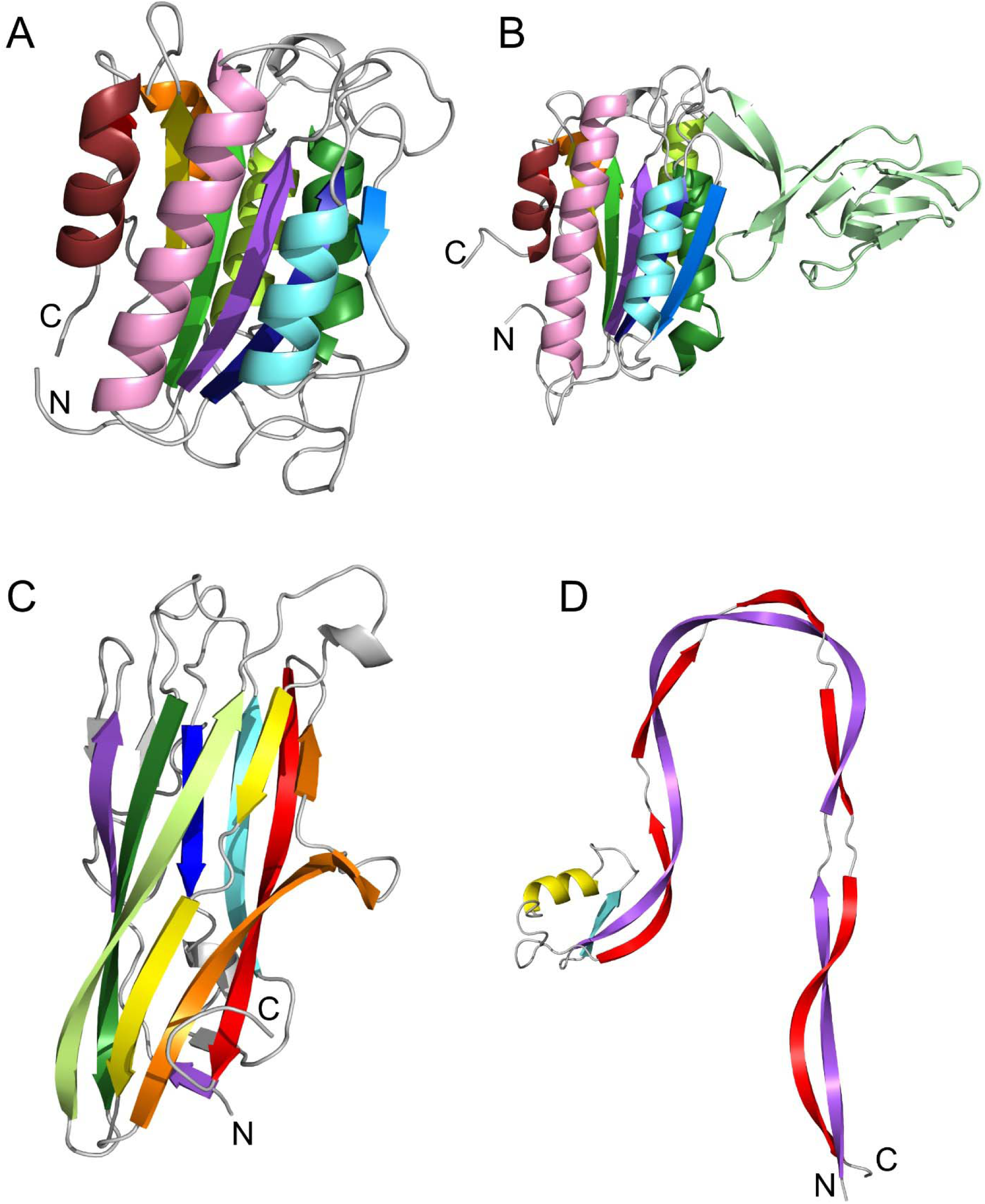
Cartoon depictions of AlphaFold3 models of various LBDs. A) vWFA domain of *Lp*1a. B) vWFA domain with insert from *As*1a. C) PDB domain from *Vv*1e. D) Unknown domain from *Ab*1. The twelve Β strands and α helices in vWFA domains are coloured from N to C terminus as follows: purple strand, pink helix, dark blue strand, medium blue strand, cyan helix, forest green helix, green strand, yellow-green helix, yellow strand, orange helix, red strand, dark red helix. The unknown domain that buds off between the green strand and yellow-green helix in B is coloured pale green. The ten β strands in the PBD are coloured in the same order but green and dark red are omitted. The unknown domain in D is colored purple, cyan, yellow and red from N to C terminus.

**Supplementary Table 1.**
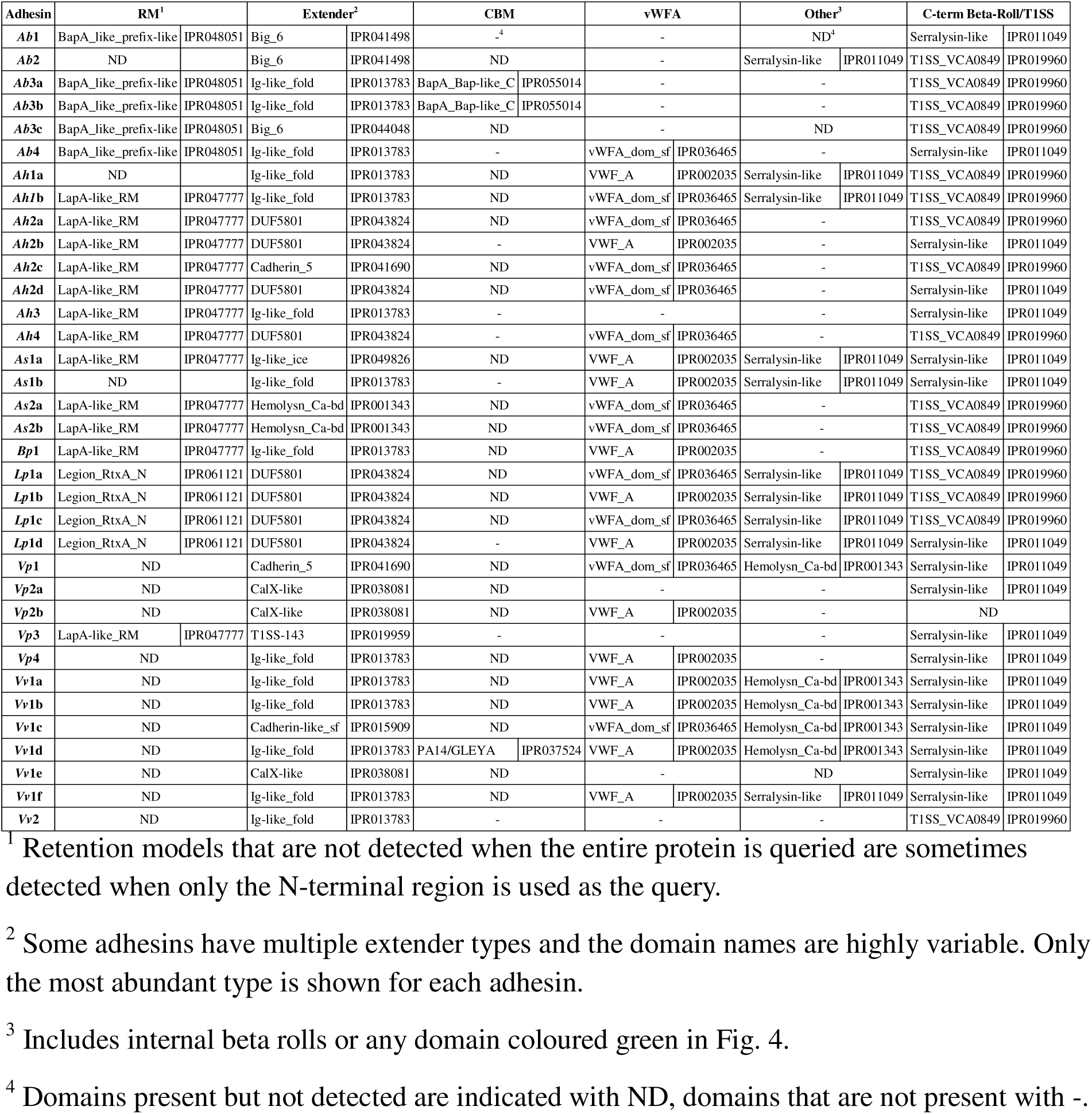
Short names and InterPro accession numbers for domains detected by InterProScan ^42^ in each isoform.

**Supplementary Table 2.** Details of gene status at each adhesin locus.

| Locus<sup>2</sup> | Adhesin | Count | Intact Genes | Missannotated | Unique frameshift | Shared frameshift | TE insertion | Absent | # of high identity repeats<sup>2</sup> |
| --- | --- | --- | --- | --- | --- | --- | --- | --- | --- |
| 1 | *Ab1* | 38 | 34 | 0 | 3 | 0 | 1 | 12 | 6 |
| 2 | *Ab2* | 29 | 28 | 0 | 1 | 0 | 0 | 21 | 2 |
| 3 | *Ab3a* | 42 | 14 | 0 | 3 | 25 | 0 | 0 | 22 |
|  | *Ab3b* | 5 | 5 | 0 | 0 | 0 | 0 |  | 28 |
|  | *Ab3c* | 3 | 2 | 0 | 1 | 0 | 0 |  | 13 |
| 4 | *Ab4* | 3 | 3 | 0 | 0 | 0 | 0 | 47 | 18 |
| 1s1 | *Ah1a* | 42 | 27 | 0 | 15 | 0 | 0 | 0 | 22 |
|  | *Ah1b* | 1 | 0 | 0 | 1 | 0 | 0 |  | 48 |
| 2s2 | *Ah2a* | 24 | 1 | 19 | 3 | 0 | 1 | 0 | 13 |
|  | *Ah2b* | 11 | 0 | 9 | 1 | 0 | 1 |  | 12 |
|  | *Ah2c* | 6 | 5 | 0 | 1 | 0 | 0 |  | 6 |
|  | *Ah2d* | 2 | 1 | 1 | 0 | 0 | 0 |  | 14 |
| 3 | *Ah3* | 28 | 26 | 0 | 2 | 0 | 0 | 15 | 0 |
| 4 | *Ah4* | 1 | 0 | 1 | 0 | 0 | 0 | 42 | 29 |
| 1s1 | *As1a* | 35 | 28 | 0 | 6 | 0 | 1 | 0 | 8 |
|  | *As1b* | 1 | 1 | 0 | 0 | 0 | 0 |  | 65 |
| 2s2 | *As2a* | 31 | 20 | 1 | 1 | 9 | 0 | 0 | 0 |
|  | *As2b* | 5 | 2 | 0 | 3 | 0 | 0 |  | 7 |
| 1 | *Bp1* | 50 | 50 | 0 | 0 | 0 | 0 | 0 | 3 |
| 1 | *Lp1a* | 6 | 6 | 0 | 0 | 0 | 0 | 0 | 14 |
|  | *Lp1b* | 21 | 16 | 0 | 5 | 0 | 0 |  | 23 |
|  | *Lp1c* | 1 | 0 | 0 | 0 | 0 | 0 |  | 20 |
|  | *Lp1d* | 22 | 21 | 0 | 1 | 0 | 0 |  | 25 |
| 1 | *Vp1* | 50 | 20 | 28 | 2 | 0 | 0 | 0 | 5 |
| 2 | *Vp2a* | 46 | 45 | 0 | 1 | 0 | 0 | 0 | 13 |
|  | *Vp2b* | 4 | 4 | 0 | 0 | 0 | 0 |  | 18 |
| 3 | *Vp3* | 50 | 48 | 0 | 2 | 0 | 0 | 0 | 5 |
| 4 | *Vp4* | 1 | 1 | 0 | 0 | 0 | 0 | 49 | 11 |
| 1 | *Vv1a* | 10 | 9 | 0 | 1 | 0 | 0 | 0 | 7 |
|  | *Vv1b* | 7 | 4 | 0 | 3 | 0 | 0 |  | 54 |
|  | *Vv1c* | 7 | 3 | 0 | 4 | 0 | 0 |  | 29 |
|  | *Vv1d* | 4 | 2 | 0 | 2 | 0 | 0 |  | 50 |
|  | *Vv1e* | 1 | 1 | 0 | 0 | 0 | 0 |  | 15 |
|  | *Vv1f* | 1 | 0 | 0 | 1 | 0 | 0 |  | ~67 |
| 2 | *Vv2* | 29 | 29 | 0 | 0 | 0 | 0 | 1 | 0 |

<sup>1</sup> Loci are numbered as in Fig. 5 with those that are syntenic indicated with the same s#.
<sup>2</sup> Repeat number varies between isoforms, count given for the isoform listed in Table 3.

**Supplementary Table 3.**
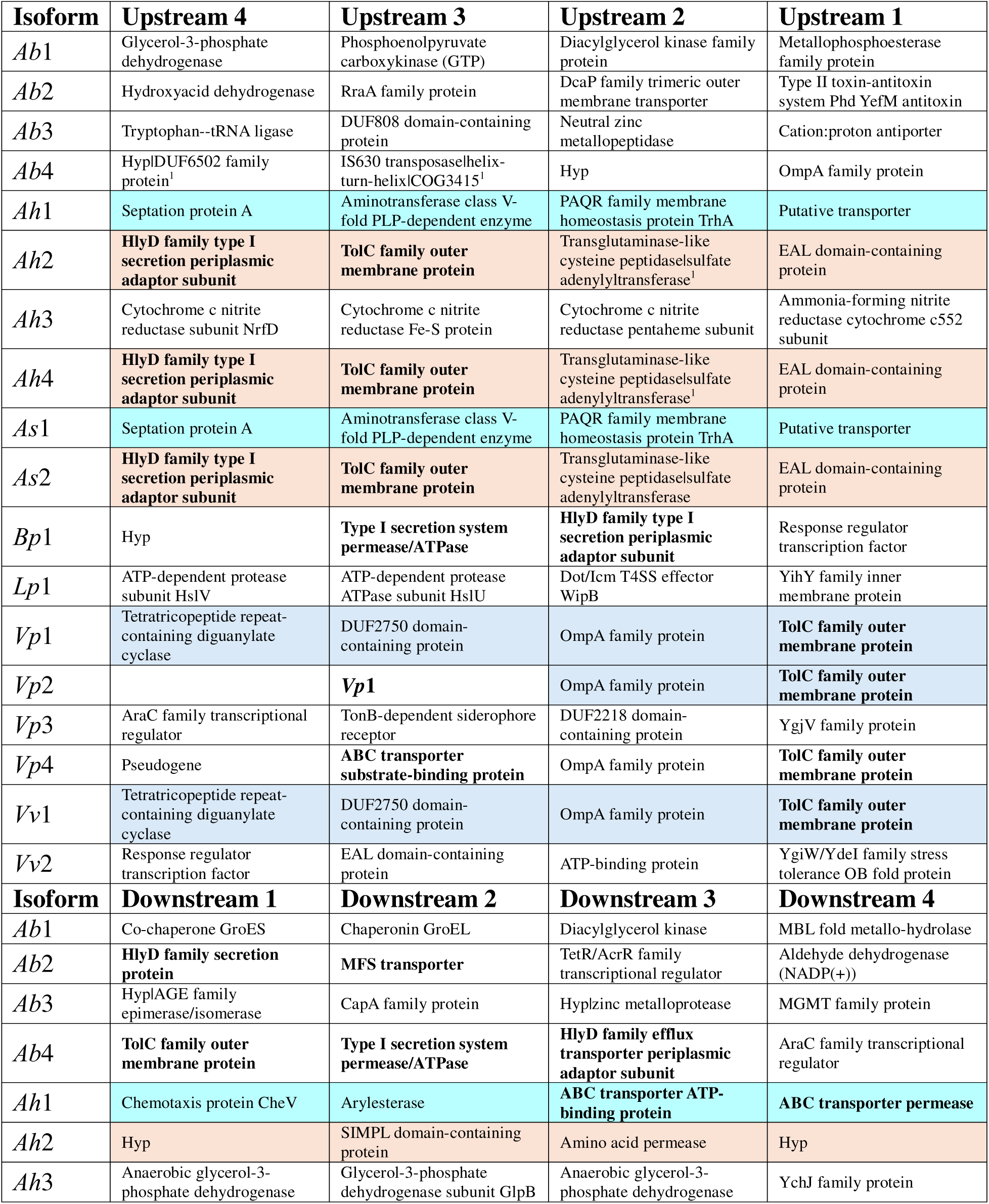
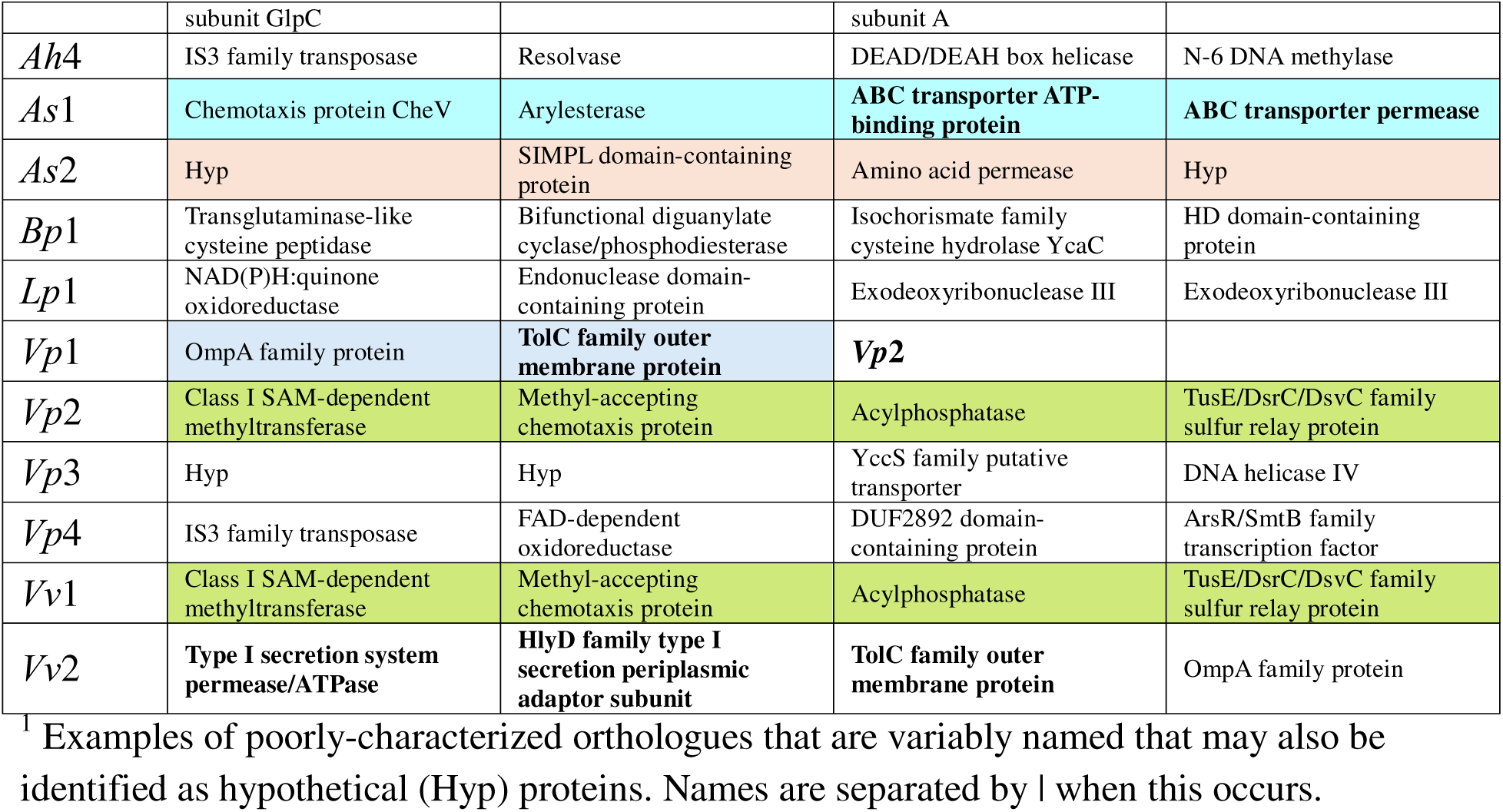
Names of the proteins encoded by the four genes found immediately upstream and downstream of each adhesin locus. Shading indicates shared microsynteny and matches the colouring used in Fig. 5. Components of the type 1 secretion system are bolded.

**Supplementary Table 4.** Frameshifts and sequencing type for A. hydrophila locus 1a.

| Isoform | Frameshifted (%) |  | Intact (%) |  | Number<br>frameshifted | Number<br>Intact |
| --- | --- | --- | --- | --- | --- | --- |
|  | PacBio | Nanopore | PacBio | Nanopore |  |  |
| *Ah1a* | 80 | 20 | 30 | 70 | 15 | 27 |

